# Histopathological Landscape of Molecular Genetics and Clinical Determinants in MDS Patients

**DOI:** 10.1101/2020.05.03.073858

**Authors:** Oscar Brück, Susanna Lallukka-Brück, Helena Hohtari, Aleksandr Ianevski, Freja Ebeling, Panu Kovanen, Soili Kytölä, Tero Aittokallio, Pedro Marques Ramos, Kimmo Porkka, Satu Mustjoki

**Author notes:** Co-corresponding authors Drs S Mustjoki and O Brück, Hematology Research Unit Helsinki, University of Helsinki and Helsinki University Hospital Comprehensive Cancer Center, Haartmaninkatu 8, Helsinki FIN-00290, Finland. Equal contribution (shared 2^nd^ authors) (Susanna Lallukka-Brück, Helena Hohtari, Aleksandr Ianevski).

## Abstract

In myelodysplastic syndrome (MDS), bone marrow (BM) histopathology is visually assessed to identify dysplastic cellular morphology, cellularity, and blast excess. Yet, many morphological findings elude the human eye. Here, we extracted visual features of 236 MDS, 87 MDS/MPN, and 10 control BM biopsies with convolutional neural networks. Unsupervised analysis distinguished underlying correlations between tissue composition, leukocyte metrics, and clinical characteristics. We applied morphological features in elastic net-regularized regression models to predict genetic and cytogenetic aberrations, prognosis, and clinical variables. By parallelizing tile, pixel, and leukocyte-level image analysis, we deconvoluted each model to texture and cellular composition to dissect their pathobiological context. Model-based mutation predictions correlated with variant allele frequency and number of affected genes per pathway, demonstrating the models’ ability to identify relevant visual patterns. In summary, this study highlights the potential of deep histopathology in hematology by unveiling the fundamental association of BM morphology with genetic and clinical determinants.

## Introduction

Current diagnosis of myelodysplastic syndrome (MDS) is based on identifying cellular dysplasia by visual inspection of bone marrow (BM) aspirate or biopsy.^1^ Karyotype status, blast proportion, and peripheral blood (PB) cell count are assessed for disease subclassification, according to WHO guidelines and for risk stratification by the Revised International Prognostic Scoring System (IPSS-R) criteria.^1,2^

Deep learning enables accurate visual pattern recognition with convolutional neural networks (CNN), where multiple processing layers detect and nonlinearly deconvolute image data into activation vectors.^3^ CNNs have recently led to significant breakthroughs in the analysis of biomedical images, such as diagnosis of skin tumors, retinal disease, intracranial hemorrhage, and breast cancer.^3–8^ In the context of routine hematoxylin and eosin (H&E) tissue stains, similar algorithms have improved Gleason scoring in prostate cancer, outcome prediction in colorectal cancer, and even discrimination of solid cancer patients by driver mutation status.^5,9–11^

Here, we investigate the potential of CNN-based morphological analysis in hematology. To improve our understanding of MDS histopathology and its association with clinical factors, we predict diagnosis, prognosis, IPSS-R risk score, mutated genes, cytogenetics, and patient age and gender by utilizing solely BM morphological features. We demonstrate highest detection accuracy for point mutations, such as *TET2* and *ASXL1*, which correlate with variant allele frequency (VAF), confirming the identification of mutation-specific features. To deconvolute the nonlinear interactions between disease determinants and BM histology, we introduce a novel multidimensional image analysis approach that combines information at tile, segmented white blood cell (WBC) and pixel levels, ultimately facilitating the interpretation of complex BM histopathological patterns.

## Results

### Unsupervised modelling of BM morphology revealed morphologic lineages and distinct myelodysplastic clusters

To dissect the multiple abstraction levels of BM morphology, we extracted ImageNet-configured visual activations of H&E-stained BM biopsies from patients with MDS (n=236), myelodysplastic/myeloproliferative neoplasms (MDS-MPN) (n=87) and healthy control (n=10), using VGG16 and Xception CNN infrastructures (Fig. 1a). Images of tissue microarray (TMA) cores were grayscaled and split into 500 tiles to amplify robustness and granularity. To investigate the spectrum of CNN texture patterns, we mapped image tiles from diagnostic BM samples of MDS, MDS/MPN, and control subjects with two-dimensional uniform manifold approximation and projection (UMAP) representation (Fig. 1b). Unsupervised image segregation was principally driven by stromal and cellular texture. Images with cellular content were observed to subcluster according to WBC and red blood cell (RBC) abundance, and lipid droplet density. As expected, MDS/MPN patients harbored an increased number of hypercellular tiles, while MDS patients demonstrated heterogeneous histopathological phenotypes (Fig. 1b). Notably, tiles from healthy subjects represented a harmonized balance of cellularity and lipid droplets with scarce stroma.

**Fig. 1.**
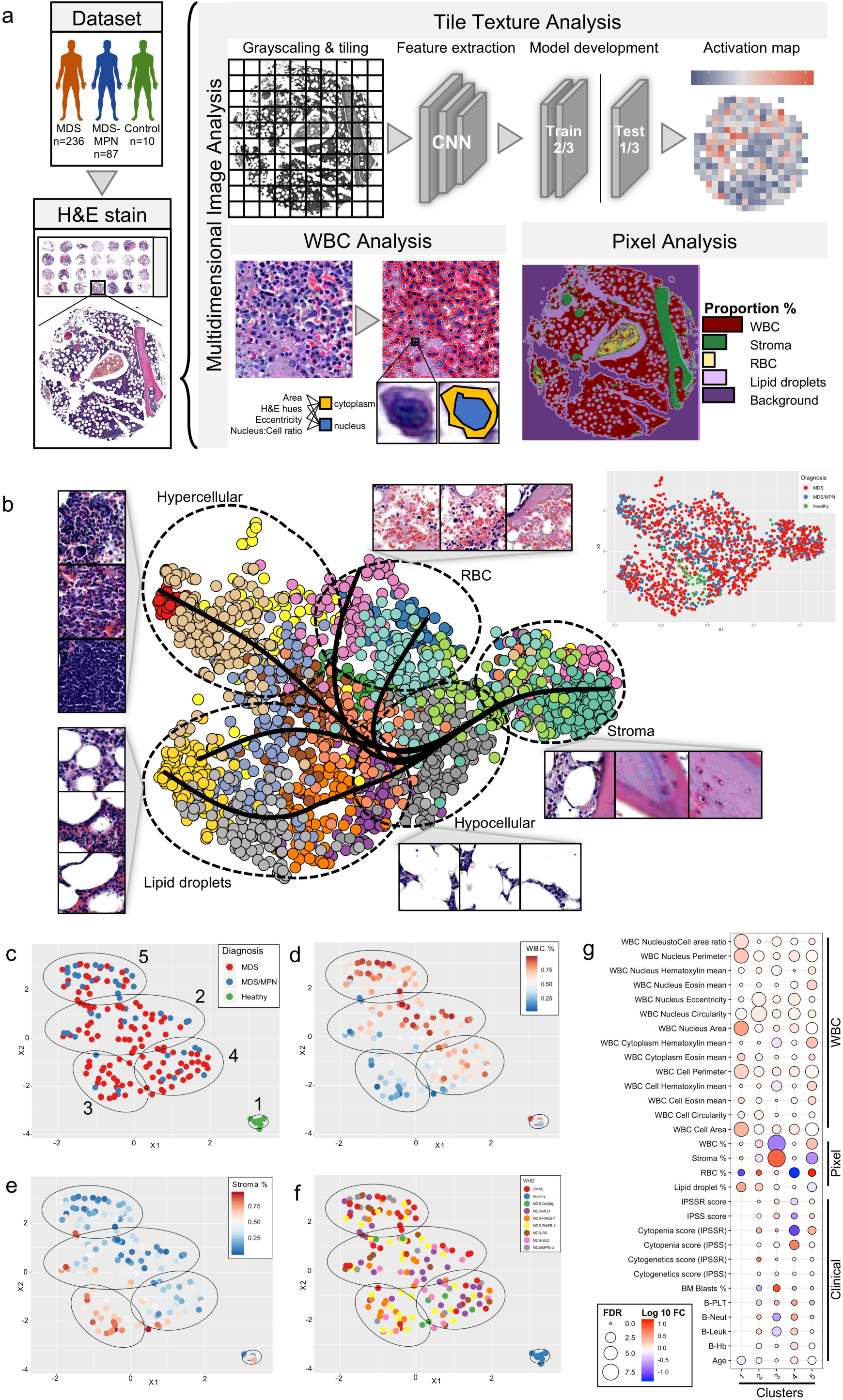
Study design. (a) Tissue microarrays (TMA) were constructed from formalin-fixed paraffin-embedded bone marrow (BM) trephine biopsies stained for hematoxylin and eosin. Images were analyzed at tile, pixel, and white blood cell-level (WBC). In the tile-level analysis, TMA spot images were split into patches, and morphological features were extracted with ImageNet-pretrained convolutional neural networks. We developed elastic net -regularized algorithms to predict multiple clinical and molecular genetics variable using only visual feature activations. Tile-level feature activations were visually merged as activation maps to deconvolve spatial prediction probability. A Weka pixel classifier was trained to identify WBC, red blood cell (RBC), stroma, and lipid droplet from images. Moreover, WBCs were segmented, and morphological metrics extracted. (b) Uniform manifold approximation and projection (UMAP) of image tile feature vectors from diagnostic myelodysplastic syndrome (MDS), myelodysplastic/myeloproliferative neoplasm syndrome (MDS/MPN), and healthy subjects. Each patient sample is presented here with random 10 tiles. Colors are encoded according to phenograph cluster (larger image) or diagnosis (right-most image). Clusters have been grouped into morphological subgroups (dashed circles). Result of a slingshot analysis has been superposed to demonstrate visual feature trajectories. (c) UMAP projection feature vectors mean-aggregated at the sample-level, clustered with k-means and color-labelled by the corresponding diagnose, (d) tissue proportion of WBC and (e) stroma, and (f) diagnostic WHO classification. (g) Each k-means cluster has been compared to remaining clusters (Wilcoxon test and Benjamin&Hochberg p-value correction) for segmented WBC and pixel-level image analysis parameters, and clinical information. Clinical information is not reported for healthy patients (Cluster 1).

PhenoGraph-driven clustering of image tiles resulted in detailed tissue patterns, which we hypothesized to form dynamic morphological entities (Fig. 1b). Slingshot lineage analysis demonstrated potential developmental trajectories between histopathological phenotypes such as diffuse transition from hypocellular to lipid droplet-dense areas^12^. Missing intercluster connections between high cellularity and lipid droplet-dense morphologies were also revealed confirming mutually exclusive pathologies.

Similar texture classes were also discovered when studying MDS image tiles from diagnosis and follow-up (Extended Data Fig. 1a). WBC proportion was reduced in follow-up samples compared to baseline but tended to recover in later timepoints reflecting transition from frequent disease-modifying treatments at early follow-up and blast expansion at later timepoints (Extended Data Fig. 1b-d). In contrast, the proportion of stroma increased in follow-up samples. Interestingly, lower lipid droplet density was measured in follow-up samples and its level further decreased in later timepoints in samples with low blast burden. No difference in RBC levels by time or blast burden was observed.

Tile features from MDS, MDS/MPN, and healthy subjects were averaged at the sample level and 2D-projected with UMAP (Fig. 1c and http://hruh-20.it.helsinki.fi/mds_visualization). Unsupervised k-means clustering was utilized for subgrouping aggregated vectors. A distinct cluster representing healthy subjects (Cluster 1) and four myelodysplastic clusters were principally formed according to tissue texture content including the proportion of WBC, stroma, and lipid droplets and RBC and WBC metrics (Fig. 1d-g).

Interestingly, the unsupervised clustering structure of MDS and MDS/MPN samples was in line with the World Health Organization (WHO) disease classification (Fig. 1f). Cluster 3, defined histologically by high bone stroma content, was enriched with elevated blast type 1 (EB-1) and 2 (EB-2) MDS subtypes, dimmer hematoxylin staining, and leukopenic PB blood count (Fig. 1g). Cluster 5 was characterized with high WBC proportion and hypercellular BM typical for chronic myelomonocytic leukemia (CMML) and unclassifiable MDS/MPN-U patients. Moreover, these were also associated with darker hematoxylin staining and higher IPSS-R cytopenia score. Cluster 4 harbored histologically hypoplastic MDS, WHO-classified as single-lineage or multilineage dysplasia MDS as well as low erythroid frequency linked with Del(5q) MDS^13^. The remaining cluster 2 was characterized by increased cellularity.

### The MDS BM morphology is linked to mutation, karyotype, gender, and prognostic status

MDS is characterized by recurrent oncogenic somatic variants in driver genes and chromosomal aberrations (Fig. 2a-b)^14–16^. We adapted a transfer learning approach, where elastic net-regularized regression was developed with VGG16 and Xception network feature activations extracted from tile-level H&E images. Following sample-level prediction average-pooling, we observed high inference notably for *TET2, ASXL1*, and *STAG2* mutations, chromosome 7 monosomy, and 7q deletion (Fig. 2c-f).

**Fig. 2.**
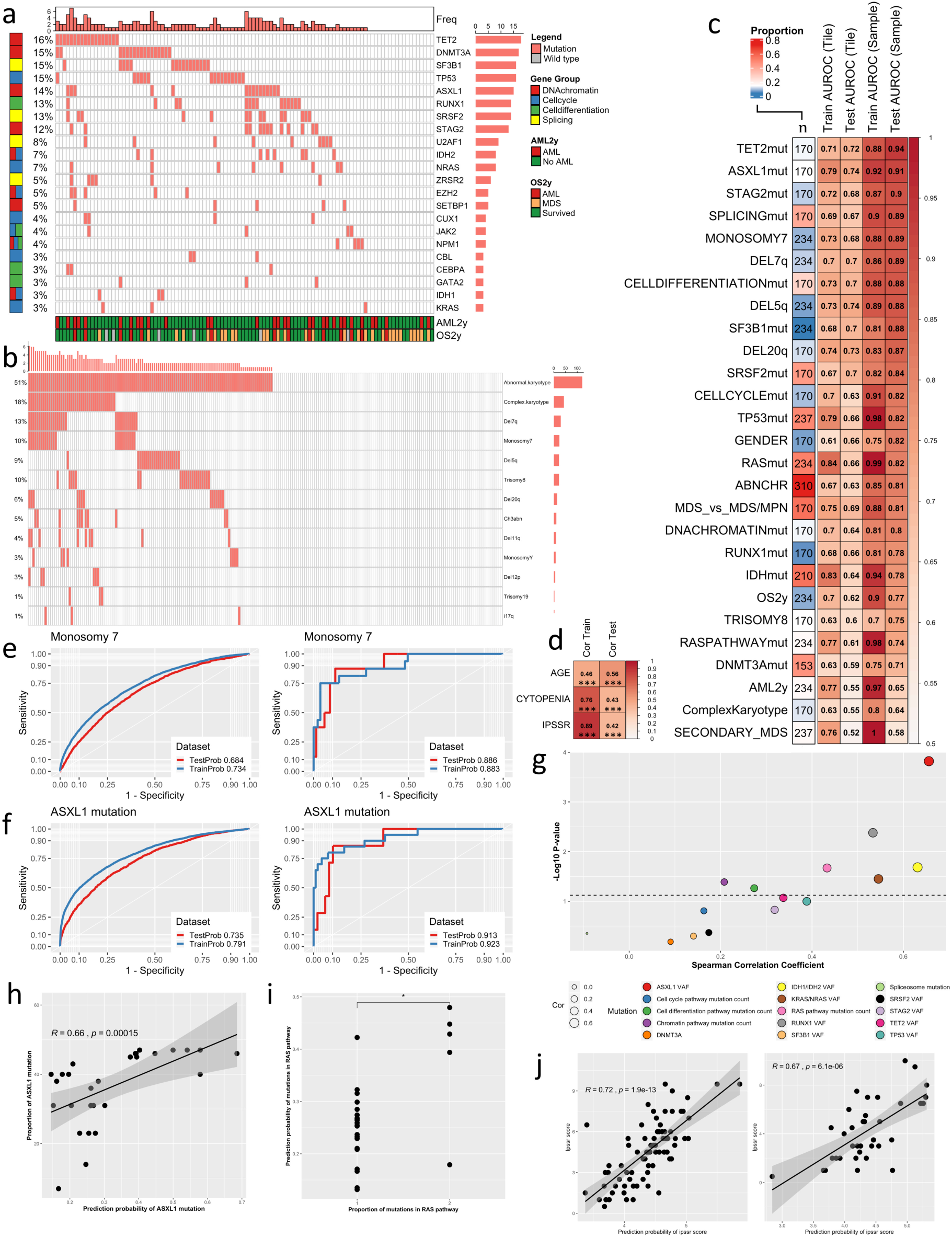
Supervised learning on bone marrow (BM) morphological features. (a) Oncoprint visualization of mutation pattern and gene groups in myelodysplastic syndrome (MDS) patients. (b) Oncoprint visualization of cytogenetics pattern in samples from MDS patients. (c) Heatmap displaying area under the receiver operating characteristic curve (AUROC) values of elastic net -regularized logistic regression models. The left-most column shows the number of samples included in the analysis and the cell color represent the distribution of the binary variables to be predicted. The following two columns inform the AUROC values of the models in the training (2/3) and test (1/3) dataset in the tile-level images and the last two columns at the tissue microarray (TMA) spot-level. (d) Similar plot for elastic net -regularized linear regression models. The left-most column shows the Spearman correlation value in the training dataset and right-most in the test dataset. (e) Tile-level (left) and TMA spot-level (right) AUROC for the logistic regression of monosomy 7 and (f) ASXL1 mutation status. (g) Scatter plot for the Spearman correlation (x-axis) between logistic regression predicting mutation status and the observed gene variant allele frequency for individual genes or number of genes mutated for functional pathways. (h) Linear regression (R represents Spearman correlation) between prediction probability of ASXL1 mutation and its detected variant allele frequency. (i) Wilcoxon comparison for predicted mutation probability and detected frequency of altered genes in the *RAS* pathway. (j) Linear regression (R represents Spearman correlation) between the predicted and observed IPSS-R score in the training (left) and test dataset (right).

Moreover, morphological features were associated with point mutations in genes regulating splicing, cell differentiation, and cell cycle, which are commonly affected in MDS. Activations extracted with the Xception infrastructure were observed to be more generalizable compared to VGG16, concomitant with higher feature heterogeneity observed in Xception correlation matrix (Extended Data Fig. 2-3). In addition, lasso and elastic net penalization provided more generalizable models than ridge regression.

Next, we investigated the validity of models predicting mutated genes and dysregulated pathways. Notably, the inferred probability of a distinct mutation correlated significantly with its variant allele frequency for *ASXL1, KRAS/NRAS, IDH1/IDH2*, and *RUNX1* genes, as well as trended for *TET2* and *TP53* genes (Fig. 2g-h and Extended Data Fig. 4). Moreover, the predicted likelihood of *RAS*, cell differentiation, and chromatin structure regulating gene pathway dysregulation was noted to correlate with the number of genes mutated in the respective pathways (Fig. 2i). Of note, the initial models were developed with all patients with mutation data, while the correlation analysis was restricted to samples with known mutation, indicating BM tissue morphology to be impacted by molecular genetics and emphasizing the algorithms’ ability to identify variant-related histopathological patterns in an unprecedented fashion.

Contrary to histopathological routines of solid tumors, tissue morphology is unutilized in risk stratification of MDS patients. Instead, the IPSS-R score accounting for PB cell count, BM blast burden, and cytogenetics is the most-established stratification tool, and associated here with AML risk and OS (Extended Data Fig. 5a-b)^2^. Of note, we could predict IPSS-R score, 2-year OS and progression to AML by solely employing H&E-stained slides (Fig. 2c-d, 2j). Moreover, deep BM morphology was associated with higher concordance of progression to AML and OS in 2 years than IPSS-R score either individually or combined with deep histopathology (Extended Data Fig. 5c-d). These results might be impacted by differences in patient selection, as our cohort represented an unbiased real-world cohort, while patients treated with disease-modifying therapies were excluded from the landmark prognostic study by Greenberg *et al*.^2^ However, patient prognostication and treatment stratification could be improved by including morphological features.

### Deconvolution of supervised prediction models with multilevel image analysis

In clinical practice, MDS is differentially diagnosed from MDS/MPN by visual inspection of cytomorphology and histopathology, evaluation of PB counts, flow cytometry, karyotype, and increasingly, molecular genetics. Tissue samples were partitioned into small TMA cores possibly limiting effective diagnostic segregation. Yet, we could discern MDS patients from MDS/MPN patients with an AUROC validation accuracy of 0.81 (Fig. 2c). To dissect the prediction model, we combined tile-level tissue texture predictions with pixel and WBC-level metrics (Fig. 3a, Extended Data Fig. 6). Pixel classification was trained to identify WBCs, RBCs, stroma including fibrotic stroma and bone trabeculae, as well as lipid droplets, composing the major tissue elements in standard H&E staining. Additionally, WBCs were segmented to extract nuclear, cytoplasmic, and pancellular measurements such as size, circularity, and hematoxylin and eosin dye variations. As expected, MDS likelihood increased if the sample represented hypoplastic texture (Fig. 3b-c). MDS morphology was also associated with dimmer hematoxylin and eosin staining and larger cell size likely due to technically larger cell segmentation area in hypocellular BM (Extended Data Fig. 7a-d).

**Fig. 3.**
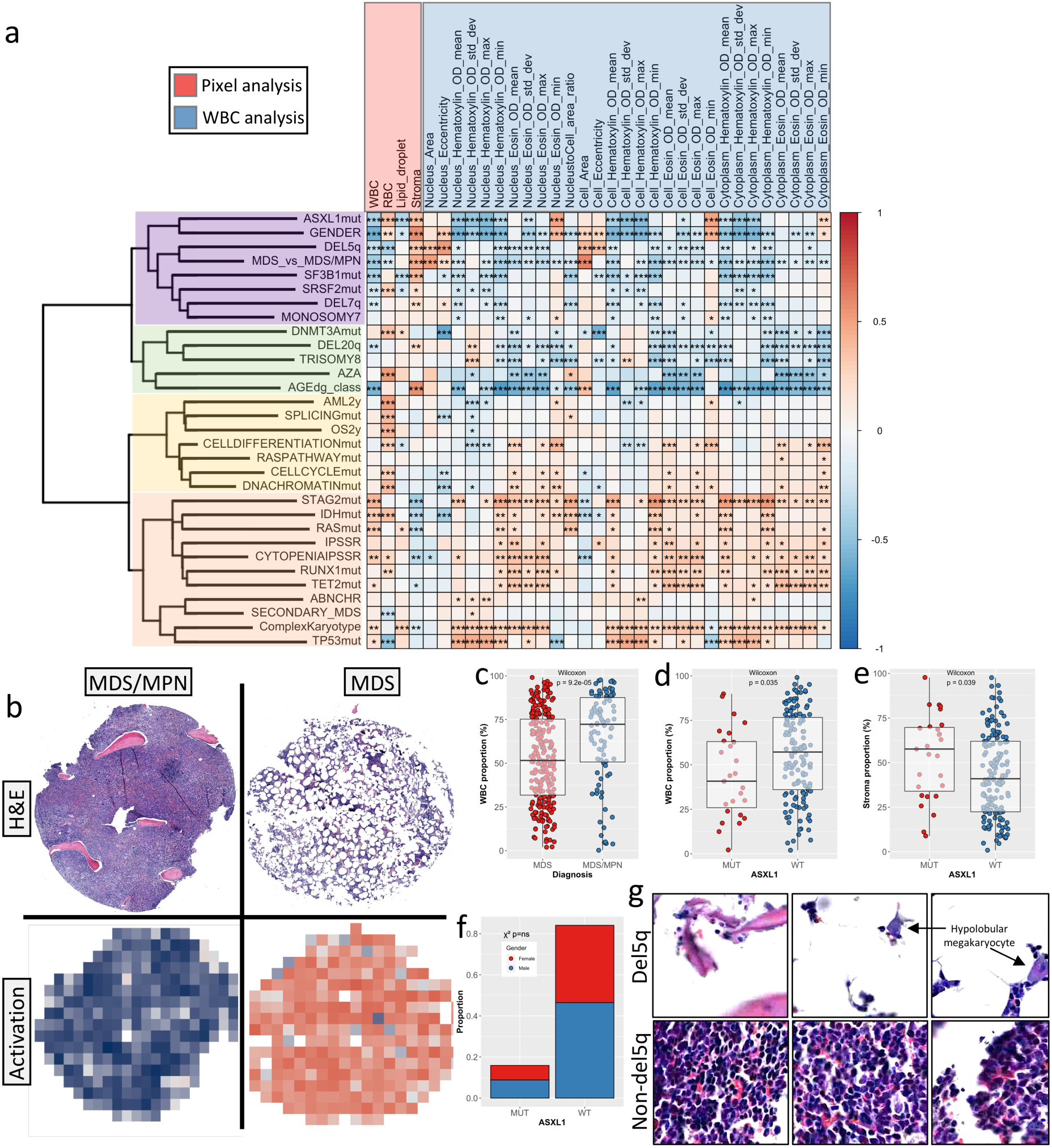
Deconvolution of supervised prediction models. (a) Correlation matrix for logistic and linear regression model predictions (rows) and pixel-level and segmented white blood cell-level (WBC) image analysis metrics aggregated per sample. The color of individual matrix cells represents the Spearman correlation and the asterisks the Benjamin-Hochberg-adjusted significance values: * p<0.05, ** p<0.01, *** p<0.001. (b) H&E-stained TMA spots and corresponding activation maps for the prediction of myelodysplastic syndrome or myelodysplastic/myeloproliferative neoplasm diagnosis (MDS or MDS/MPN). For the activation map, tile-level predictions haves been color-scaled (blue: high probability for MDS/MPN, red: high probability for MDS). (c) Scatter plot and Wilcoxon test to compare WBC proportion by MDS and MDS/MPN diagnoses. Boxplots define the interquartile ranges and median values by diagnoses. (d) Scatter plot and Wilcoxon test to compare WBC proportion and (e) stroma proportion by *ASXL1* mutation status. Boxplots define the interquartile ranges and median values by diagnoses. (f) Barplot comparing gender distribution by ASXL1 mutation status. Gender frequencies have been compared with Chi^2^ test. (g) Image tiles representing highest and lowest computed probability of chromosome 5q deletion.

Following model deconvolution, *ASXL1* mutation probability was observed to correlate with elevated accumulation of stroma and RBCs, depletion of WBCs and lipid droplets as well as reduced intracellular hematoxylin staining (Fig. 3a, Extended Data Fig. 6b). We confirmed the association between sequenced *ASXL1* mutation and lower WBC and higher stroma content from clinical data (Fig. 3d-e). Interestingly, covariates involved in predicting *ASXL1* mutation coincided with female gender inference, although *ASXL1* mutations were observed to divide equally between genders (Fig. 3f).

Chromosome 5q deletion has been described to associate with a decrease in erythroid precursor cells and increase in hypolobular megakaryocytes^13^. While RBCs have been commonly interpreted as tissue processing artefact, we observed a clear correlation between 5q deletion probability and infrequent RBCs (Fig. 3a and 3g and Extended Data Fig. 6c). In addition, when inspecting tiles associated with highest likelihood for 5q deletion, we discovered enrichment of megakaryocytes with abnormally circular nuclei (Fig. 3g).

We also noted a distinct cluster (yellow-colored in Fig. 3a) consisting of likelihood of gene pathway dysregulation, progression to AML, and OS. While we could accurately detect spliceosome (validation AUROC 0.89) and cell differentiation mutations (validation AUROC 0.88), these models were challenging to deconvolute implying association with heterogenous and complex tissue texture determinants.

## Discussion

Here, we demonstrate how the intricate and heterogeneous BM morphological landscape can be decomposed and associated with clinical data using multilevel computer vision. Remarkably, highest prediction accuracy of deep BM morphology was noted for mutation and cytogenetic aberrations, which even outweighed reported inference in solid tumors^5,9,17,18^. We suspect homogenous BM tissue consistency and lower mutation burden of MDS to account for the improved results.

The black box dilemma hinders clinical translation of deep learning algorithms.^19,20^ To increase model transparency, we deconvoluted CNN-extracted morphological patterns associated with molecular and clinical determinants by linking image analysis at tile, pixel, and cellular levels. We emphasize that similar holistic approaches should be further explored with neural network-based semantic and instance segmentation methods to improve dissection of complex models.

Taken together, deep mining of the BM tissue texture in a larger scale could assist pathologists by revealing intricate morphological patterns defining disease subtypes and eventually improving clinical stratification of MDS patients.

## Materials and methods

### Patients

The study population comprised MDS (n=142) and MDS/MPN patients (n=51) and control subjects (n=10) treated between 2000-2018 at the Department of Hematology in the Helsinki University Hospital (Supplementary Table 1). BM trephine biopsies taken at diagnosis and follow-up were applied for from the Helsinki Biobank. According to Helsinki University Hospital’s ethical board guidelines, BM trephine samples were collected from subjects without diagnosis of hematologic malignancy, chronic infection, nor autoimmune disorder in six years of follow-up. Control subjects were 54.5% males and 55.4 [40.0-82.0] years old at the time of BM sampling. MDS and MDS/MPN patients were older than control subjects (p=0.02 and p=0.002, respectively) but did not differ significantly by gender distribution (Supplementary Table 1). All subjects gave written informed research consent. The study complied with the Declaration of Helsinki and the HUS ethics committee (DNRO 303/13/03/01/2011). All clinical data were collected from the HUS hematology datalake, a GDPR-compliant database integrating data from electronical health registries and the Finnish Hematology Registry.

### Sequencing

Genomic DNA was isolated from diagnostic BM samples (n=108) using the QIAsymphony DSP DNA kit. Driver gene mutations were defined using a clinical-grade, myeloid amplicon sequencing panel capable of identifying mutations with variant allele frequency >2% (mean sequencing depth 6000x; Extended Data Fig. 8). In addition, data from additional 40 samples analyzed with the Illumina TruSight myeloid amplicon sequencing panel were included (mean sequencing depth 100x).

### Tissue Microarrays (TMAs)

Upon sampling, fresh BM biopsies were conformed to routine formalin-fixation and paraffin-embedding (FFPE). TMAs were cast by a single 2 mm core (MDS and MDS/MPN) or double 1 mm cores (controls) per sample from representative BM biopsy areas (Fig. 1a). Tissue blocks were cut into 4 μm thick sections and stained with hematoxylin and eosin (H&E). Slides were digitized at 0.22 μm/pixel (20x objective magnification) with the whole-slide scanner Pannoramic 250 FLASH (3DHISTECH Ltd.).

### Image preprocessing

H&E images were preprocessed by converting RGB images into grayscale to avoid bias from technical artifacts caused by sample processing and to increase the robustness of texture features. Moreover, we standardized non-tissue background pixels by converting pixels exterior to binary tissue masks into 255 (white). H&E images were resized to 6.000 pixels in horizontal length and split into 500 equally-sized tiles. Tiles with mean pixel intensity over 240 represented non-tissue background and were excluded from the analysis resulting in 73.531 tiles. Tile size was optimized to marginally outsize the largest BM lipid droplets to avoid their classification as non-tissue background.

### Feature extraction

We adapted a transfer learning approach where image tiles activations were obtained with pretrained Xception and VGG16 convolutional networks as well as the Keras deep learning framework, which have achieved high accuracy in classifying ImageNet data^21–23^. Individual tiles were resized into equal sizes (224×224 for VGG16 and 299×299 for Xception) and rescaled (0,1) for feature extraction. For each tile, a 2.048-bin feature vector was extracted at the second last fully-connected Xception network layer. To tackle the nonidentical dimensionality of the VGG16 architecture and enable unbiased network comparison, we exported features from the last layer (n=25.088) and retained only the top 2.048 features with the highest variance-to-mean ratio (Extended Data Fig. 3).

### Regression models

To predict genomic, cytogenetic, prognostic, and patient demographics, image tiles were first split at the sample level into training (2/3) and test (1/3) datasets. Models were trained with L1 (alpha = 1) and L2-penalized (alpha = 0) and elastic net-regularized (alpha = 0.5) regression models using 5-fold cross-validation^24,25^. Lambda values were optimized for each fixed mixing parameter alpha to reach minimum cross-validation error (lambda.min) and to choose the lambda at one standard error of the minimum (lambda.1se). Training occurred at the tile level, and the prediction results were assessed at tile level and after sample-wise average-aggregation also at sample level. In summary, each predicted variable was estimated based on 12 algorithms (two CNN models and three alpha and two lambda elastic net regularization parameter values).

Separate prediction models for IPSS-R score, IPSS-R cytopenia score, and age at diagnosis were developed with linear regression and using only diagnostic MDS samples. Age was transformed into <50.0, 50.0-59.9, 60.0-69.9, 70.0-79.9, and >80.0 years age categories, which ameliorated the accuracy and interpretation of results. Gender, mutations, cytogenetic aberrations, overall survival (OS) in 2 years, AML progression in 2 years, and MDS etiology were predicted with logistic regression using both diagnosis and follow-up samples (Fig. 2c). Only genes and chromosomal aberrancies present in over 9% of samples were selected (Fig. 2a-b). Disease etiology was assigned as either “de novo” or “secondary MDS”. Azacytidine response was predicted with logistic regression using samples taken 0-365 days before treatment start.

Abbreviations used for predicted variables. SPLICINGmut: spliceosome mutations. CELLDIFFERENTIATIONmut: mutation in genes regulating cell differentiation. CELLCYCLEmut: mutation in genes regulating cellcycle. DNACHROMATINmut: mutation in genes regulating DNA chromatin structure. RASmut: mutation in *NRAS* or *KRAS*. ABNCHR: presence of any abnormal chromosome. IDHmut: mutation in *IDH1* or *IDH2*. OS2y: overall survival event in 2 years of follow-up. RASPATHWAYmut: mutation in genes regulating RAS pathway. AML2y: progression to AML in 2 years of follow-up. AZA: azacytidine treatment response.

### Pixel classification

Each RGB TMA spot images were analyzed with the trainable Weka pixel classification module of Fiji using default parameters^26^. Each pixel was classified as either WBC, RBC, stroma, or lipid droplet. Stroma included fibrotic stroma and bone trabeculae. The area of individual classes was calculated as proportion to a binary tissue mask area to estimate their relative tissue area rather than absolute quantity. The tissue mask was created by converting H&E images into binary format and performing mask dilatation, empty hole fill and mask erosion steps (Extended Data Figure 9). Lipid droplets were defined as filled image holes from the initial tissue mask.

### WBC analysis

Each RGB TMA spot images were analyzed with the open-source software QuPath (v0.2.0)^27^. WBCs were segmented with the watershed cell detection and background radius 30px, median filter radius 0px, sigma 6px, minimum area 10px^2^, maximum area 10.000px^2^, threshold 0.1, max background intensity 2, and cell expansion 5px (Extended Data Figure 7). Nucleus and cytoplasm staining intensity and size as well as cell circularity metrics were extracted for individual WBC and averaged at the TMA spot level. While not quantifying identical measurements, a clear correlation between the frequency of segmented WBCs and proportion of the WBC pixel area was observed (R = 0.52, p<0.001, Spearman correlation; Extended Data Figure 10).

### Statistical Analysis

Continuous variables were compared with Wilcoxon test (unpaired, two-tailed), and correlated with Spearman’s rank correlation coefficient. Categorical variables were compared with Chi^2^ test. P-values were adjusted with Benjamini-Hochberg’s correction when necessary.^28^ Cox regression analysis (log-rank test) was used for survival analysis. Model fitness was assessed by calculating statistical significance of area under the receiver operating characteristic curves (AUROC). AUROC values for predicting AML progression, OS and IPSS-R were further compared with DeLong’s test^29^.

For unsupervised analysis, we selected uniform manifold approximation and projection (UMAP) method, over principal component analysis (PCA), as visual features follow non-normal distribution. PhenoGraph is a graph-based community detection method designed for high-resolution single-cell data analogical to visual features^30^. Therefore, single tiles were clustered with Phenograph with default settings to attain higher morphological granularity. K-means clustering was selected for sample grouping to simplify interpretation of patient grouping, and the *k* parameter was harmonized using the consensus of 30 indices based on Euclidean distance^31^. Feature extraction, regression models, and statistical analysis were performed with R 3.5.1 and packages are listed in Supplementary Table 2^32^.

### Code and data availability

Code used for data analysis are available at https://github.com/obruck/MDS_HE_IA/. Image data, activation maps and deconvolution metrics are available at https://hruh-20.it.helsinki.fi/mds_visualization.

## Acknowledgments

We are grateful to the members of the Hematology Research Unit Helsinki for discussions and technical help. We thank the Helsinki Biobank and the Digital and Molecular Pathology Unit supported by University of Helsinki and Biocenter Finland for digital microscopy services.

## Authors Contributions

Conception and design: O.B., S.M. Collection and assembly of clinical data: H.H., K.P., O.B., S.L.-B. Collection and assembly of image data: O.B., P.K. Collection and assembly of sequencing data: F.E., O.B., S.K., S.M. Image analysis and data analysis: O.B. Visualization: A.I., O.B., T.A. Data interpretation: All authors. Manuscript writing: All authors. Final approval of manuscript: All authors.

## Disclosure of Conflicts of Interest

K.P. has received honoraria and research funding from Celgene, Incyte, Novartis and Bristol-Myers Squibb. S.M. has received honoraria and research funding from Pfizer, Novartis and Bristol-Myers Squibb. H.H. has received research funding from Incyte. P.R.M. is employed by Novartis Pharmaceuticals.

## Source of funding

This study was supported by University of Helsinki, the Doctoral Programme in Biomedicine (DPBM) and personal grants (O. Brück) from Biomedicum Helsinki Foundation, Finnish medical foundations (Suomen Lääketieteen Säätiö and Finska Läkaresällskapet), research grants (S. Mustjoki) from the Cancer Foundation Finland, Sigrid Juselius Foundation, Signe and Ane Gyllenberg Foundation, Relander Foundation, State funding for university-level health research in Finland, Helsinki Institute of Life Sciences Fellow grant, and investigator-initiated research grant from Novartis.

## Notes

http://hruh-20.it.helsinki.fi/mds_visualization

### Competing Interest Statement

The authors have declared no competing interest.

