## Supplemental Data for "Histopathological Landscape of Molecular Genetics and Clinical Determinants in MDS Patients"

Supplementary Table 1. Patient characteristics at diagnosis.

| | MDS (n=142) | MDS-MPN (n=51) |
|---|---|---|
| Gender, male % | 51.0 | 70.6 |
| Age, mean [range] | 64.8 [15.5-88.6] | 67.5 [36.4-87.0] |
| Etiology, % | | |
| • De novo | 82.5 | 92.2 |
| • Secondary | 17.5 | 7.8 |
| WHO MDS classification, % | | |
| • MDS-SLD | 14.0 | |
| • MDS-MLD | 23.1 | |
| • MDS-RS | 11.2 | |
| • MDS-EB-1 | 17.5 | |
| • MDS-EB-2 | 24.5 | |
| • MDS with isolated del(5q) | 7.0 | |
| • MDS-U | 2.8 | |
| WHO MDS/MPN classification, % | | |
| • CMML | | 68.8 |
| • MDS/MPN-U | | 31.2 |
| Survival status in 2 years after diagnosis, % | | |
| • Alive | 42.0 | 15.7 |
| • Dead | 49.0 | 84.3 |
| • Not defined or censored | 9.1 | 0.0 |
| Progression to AML in 2 years after diagnosis, % | | |
| • No | 35.7 | 60.8 |
| • Yes | 29.4 | 39.2 |
| • Not defined or censored due to alloHCST | 35.0 | 0.0 |
| Azacitidine-treated within 1 year of diagnosis, % | | |
| • No | 60.1 | 66.7 |
| • Yes | 39.9 | 33.3 |
| IPSS risk class, % | | |
| • Low | 16.8 | |
| • Intermediate-1 | 39.9 | |
| • Intermediate-2 | 18.2 | |
| • High | 16.8 | |
| • Not defined | 8.4 | |
| IPSS-R risk class, % | | |
| • Very low | 11.9 | |
| • Low | 23.1 | |
| • Intermediate | 18.9 | |
| • High | 18.2 | |
| • Very high | 19.6 | |
| • Not defined | 8.4 | |
| AlloHSCT, % | | |
| • No | 81.8 | 84.3 |
| • Yes | 16.8 | 15.7 |
| • Not defined | 1.4 | 0.0 |

Abbreviations. World Health Organization (WHO); Myelodysplastic syndrome with single lineage dysplasia (MDS-SLD); MDS with multilineage dysplasia (MDS-MLD); MDS with ring sideroblasts (MDS-RS); MDS with excess blasts 1 (MDS-EB-1); MDS with excess blasts 2 (MDS-EB-2); MDS, unclassifiable (MDS-U); Chronic myelomonocytic leukemia

(CMML); Myelodysplastic/Myeloproliferative neoplasm, unclassifiable (MDS/MPN-U); Acute myeloid leukemia (AML); Allogeneic hematopoietic stem cell transplantation (alloHCST); International Prognostic Scoring System (IPSS); Revised IPSS (IPSS-R)

Supplementary Table 2. R Packages used in the analysis

R version 3.05.01 (2018-07-02)
Platform: x86_64-apple-darwin15.6.0 (64-bit)
Running under: macOS 10.14.02

| | | | | | | | |
|---|---|---|---|---|---|---|---|
| ComplexHeatmap 2.3.2 | dendextend 1.9.0 | ggdendro 0.1-20 | corrplot 0.84 | Hmisc 4.2-0 | Formula 1.2-3 | lattice 0.20-38 | ggpubr 0.2 |
| magrittr 1.5 | ROCR 1.0-7 | gplots 3.0.1 | plotROC 2.2.1 | survAUC 1.0-5 | survival 2.43-1 | pROC 1.13.0 | biglasso 1.3-7 |
| ncvreg 3.11-1 | bigmemory 4.5.33 | glmnet 2.0-16 | foreach 1.4.7 | writexl 1.0 | reshape2 1.4.3 | NbClust 3.0 | factoextra 1.0.5 |
| raster 2.8-4 | sp 1.3-1 | ggimage 0.2.4 | RColorBrewer 1.1-2 | readxl 1.1.0 | uwot 0.0.0.9010 | Matrix 1.2-15 | Rphenograph 0.99.1 |
| igraph 1.2.4.1 | forcats 0.3.0 | stringr 1.4.0 | dplyr 0.8.3 | purrr 0.3.3 | readr 1.3.1 | tidyr 1.0.0 | tibble 2.1.3 |
| ggplot2 3.2.1 | tidyverse 1.2.1 | keras 2.2.4 | snakecase 0.9.2 | rvcheck 0.1.3 | lubridate 1.7.4 | base64enc 0.1-3 | class 7.3-14 |
| circlize 0.4.8 | backports 1.1.5 | plyr 1.8.4 | lazyeval 0.2.2 | splines 3.5.1 | tfruns 1.4 | digest 0.6.23 | htmltools 0.4.0 |
| viridis 0.5.1 | magick 2.0 | gdata 2.18.0 | fansi 0.4.0 | checkmate 1.9.4 | cluster 2.0.7-1 | modelr 0.1.2 | RcppParallel 4.4.4 |
| colorspace 1.4-1 | rvest 0.3.2 | ggrepel 0.8.1 | haven 2.0.0 | xfun 0.10 | crayon 1.3.4 | jsonlite 1.6 | bigmemory.sri 0.1.3 |
| zeallot 0.1.0 | iterators 1.0.12 | glue 1.3.1 | gtable 0.3.0 | GetoptLong 0.1.8 | kernlab 0.9-27 | shape 1.4.4 | prabclus 2.2-6 |
| DEoptimR 1.0-8 | scales 1.0.0 | mvtnorm 1.0-8 | Rcpp 1.0.3 | viridisLite 0.3.0 | htmlTable 1.13.2 | clue 0.3-57 | gridGraphics 0.3-0 |
| reticulate 1.12 | foreign 0.8-71 | mclust 5.4.2 | stats4 3.5.1 | htmlwidgets 1.5.1 | httr 1.4.1 | fpc 2.1-11.1 | acepack 1.4.1 |
| modeltools 0.2-22 | pkgconfig 2.0.3 | flexmix 2.3-14 | nnet 7.3-12 | ggplotify 0.0.3 | tidyselect 0.2.5 | rlang 0.4.4 | munsell 0.5.0 |
| cellranger 1.1.0 | tools 3.5.1 | cli 2.0.0 | generics 0.0.2 | broom 0.5.2 | yaml 2.2.0 | knitr 1.25 | robustbase 0.93-3 |
| caTools 1.17.1.1 | RANN 2.6.1 | nlme 3.1-137 | whisker 0.3-2 | xml2 1.2.2 | compiler 3.5.1 | rstudioapi 0.10 | png 0.1-7 |
| stringi 1.4.3 | trimcluster 0.1-2.1 | tensorflow 1.13.1 | vctrs 0.2.1 | pillar 1.4.2 | lifecycle 0.1.0 | GlobalOptions 0.1.1 | data.table 1.12.6 |
| bitops 1.0-6 | R6 2.4.1 | latticeExtra 0.6-28 | KernSmooth 2.23-15 | gridExtra 2.3 | codetools 0.2-15 | MASS 7.3-51.1 | gtools 3.8.1 |
| assertthat 0.2.1 | rjson 0.2.20 | withr 2.1.2 | parallel 3.5.1 | diptest 0.75-7 | hms 0.5.2 | rpart 4.1-13 | |

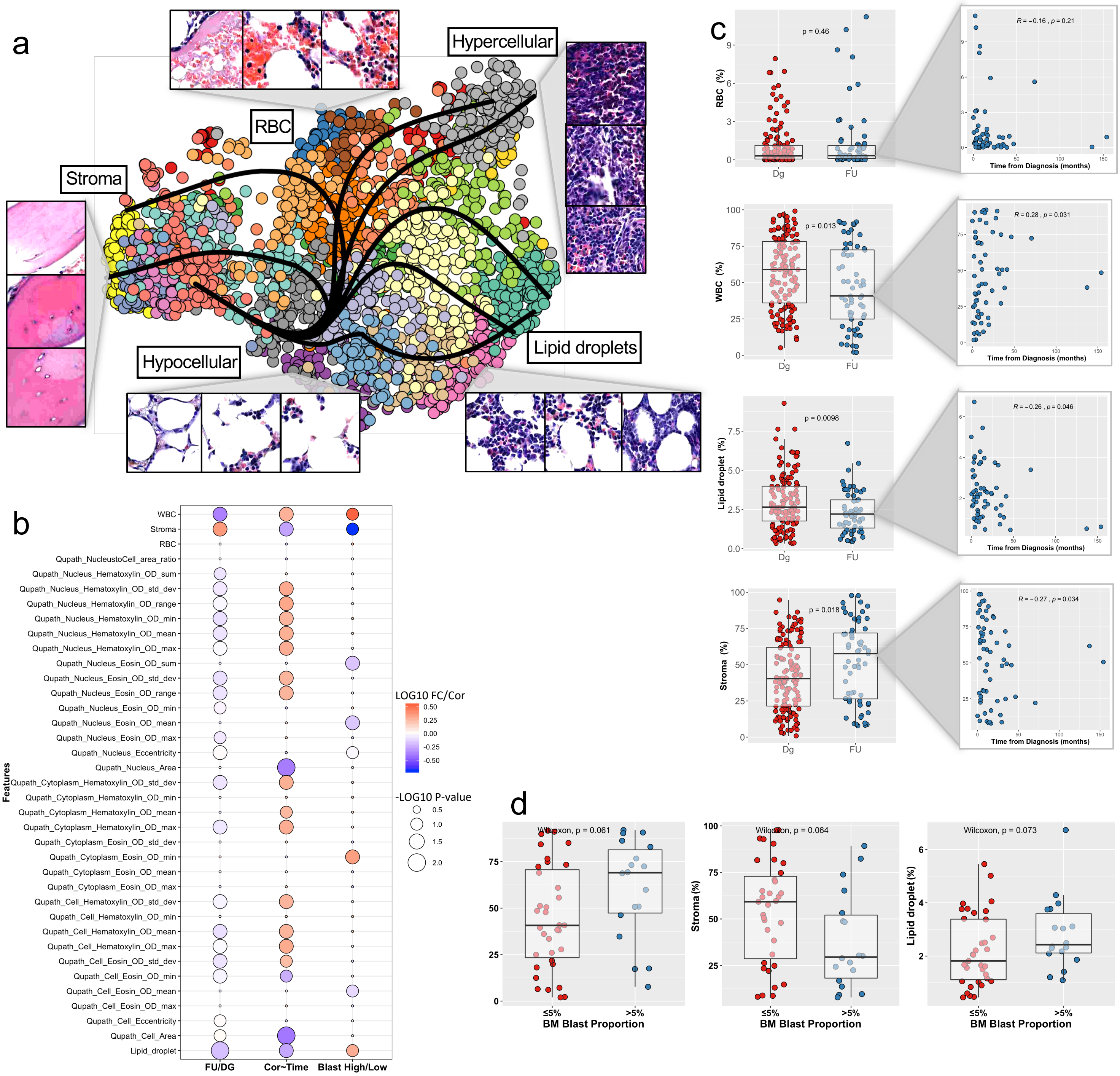


Extended Data Figure 1. (a) UMAP projection of image tiles from MDS patients at diagnosis (Dg) and follow-up (FU) clustered with PhenoGraph unsupervised analysis. Each sample is presented with random 10 tiles and their color marks their PhenoGraph cluster. Each cluster has been linked with distinctive image tiles for visualization. Slingshot analysis has been superposed to demonstrate the visual feature lineage trajectories. (b) Ballonplot summarizing results for Wilcoxon tests between follow-up and diagnose samples (FU/Dg) for multiple white blood cell and tissue texture features (FU/DG), Spearman correlation between texture features and follow-up time from diagnosis (Cor~Time) and Wilcoxon test between follow-up samples with high vs. low bone marrow blast proportion (Blast High/Low). The color scale represents the fold change in median values between groups for comparison tests (FU/DG and Blast High/Low) and correlation factor for correlation tests (Cor~Time). The circle size represents p-value and nonsignificant results have been minimized for visualization purposes. (c) Wilcoxon test for comparison of Dg and FU samples of MDS patients (left) and Spearman correlation between time from diagnosis and red blood cell (RBC), white blood cell (WBC), stroma, and lipid droplet proportion (%). The y-axis tissue texture values are quantitated with pixel classification. The boxplot represents the median and interquartile range values. Diagnostic samples have been omitted from the correlation analysis. (d) Wilcoxon test for comparison of FU samples of MDS patients between low (≤5%) and high (>5%) bone marrow blast proportion and WBC, stroma, and lipid droplet proportion (%). The y-axis tissue texture values are quantitated with pixel classification. The boxplot represents the median and interquartile range values.

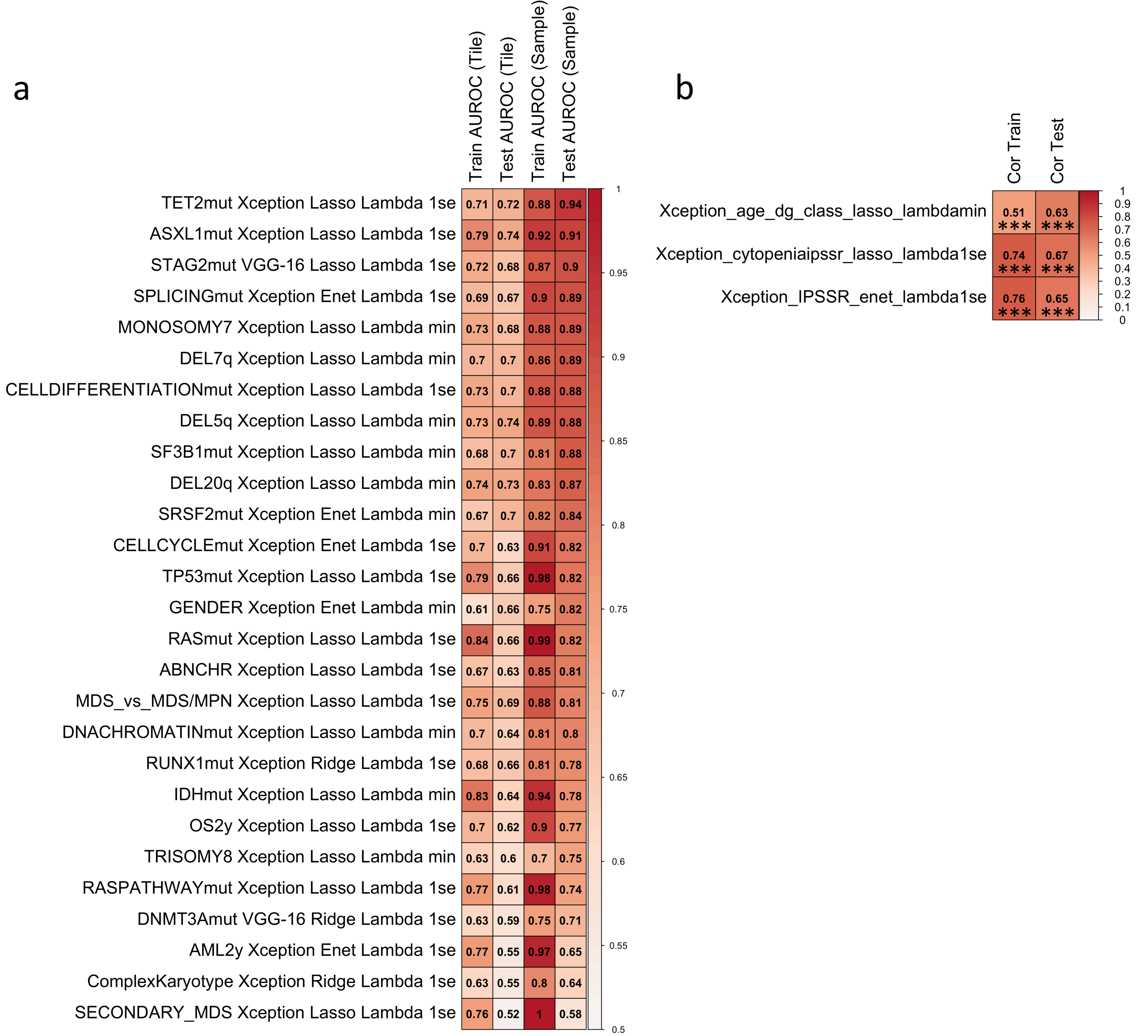


Extended Data Figure 2. Heatmap displaying area under the receiver operating characteristic curve (AUROC) values of (a) elastic net -regularized logistic regression models. Rows represent individual models and row names the predicted end point, the convolution neural network (Xception or VGG-16), elastic net alpha penalization (Lasso, Ridge or Enet) and Lambda value (Lambda min or 1se). The left-most column shows the number of samples included in the analysis. The following two columns inform the AUROC values of the models in the training (2/3) and test (1/3) dataset in the tile-level images and the last two columns at the tissue microarray (TMA) spot-level. The color grading reflects AUROC value [0.5-1.0]. (b) Similar plot for elastic net -regularized linear regression models. Rows represent individual models and row names the predicted end point, the convolution neural network (Xception or VGG-16), elastic net alpha penalization (Lasso,  Ridge or Enet) and Lambda value (Lambda min or 1se). The left-most column shows the correlation value in the training dataset and right-mostin the test dataset. The color grading reflects Spearman correlation values [0.0-1.0]. All correlations were statistically significant (*** p<0.001). Abbreviations. SPLICINGmut: spliceosome mutations. CELLDIFFERENTIATIONmut: mutation in genes regulating cell differentiation. CELLCYCLEmut: mutation in genes regulating cellcycle. RASmut: mutation in *NRAS* or *KRAS*. ABNCHR: presence of any abnormal chromosome. IDHmut: mutation in *IDH1* or *IDH2*. OS2y: overall survival event in 2 years of follow-up. RASPATHWAYmut: mutation in genes regulating RAS pathway. AML2y: progression to AML in 2 years of follow-up.

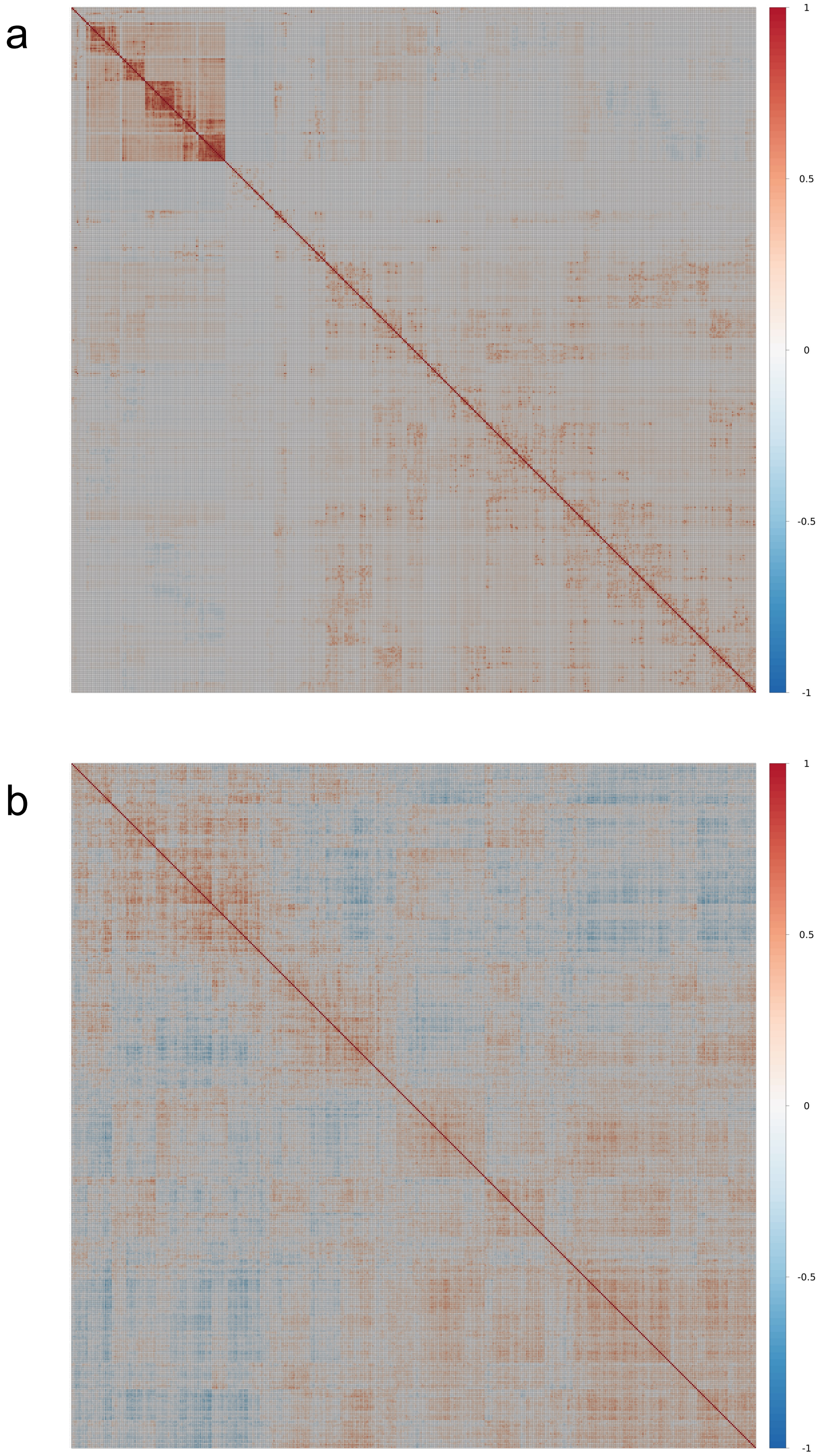


Extended Data Figure 3. Correlation plot of the visual features extracted from the (a) VGG16 and (b) Xception neural network configured with the ImageNet weights. For visualization purposes 25% (n=512) of the features used in the analysis are plotted here.

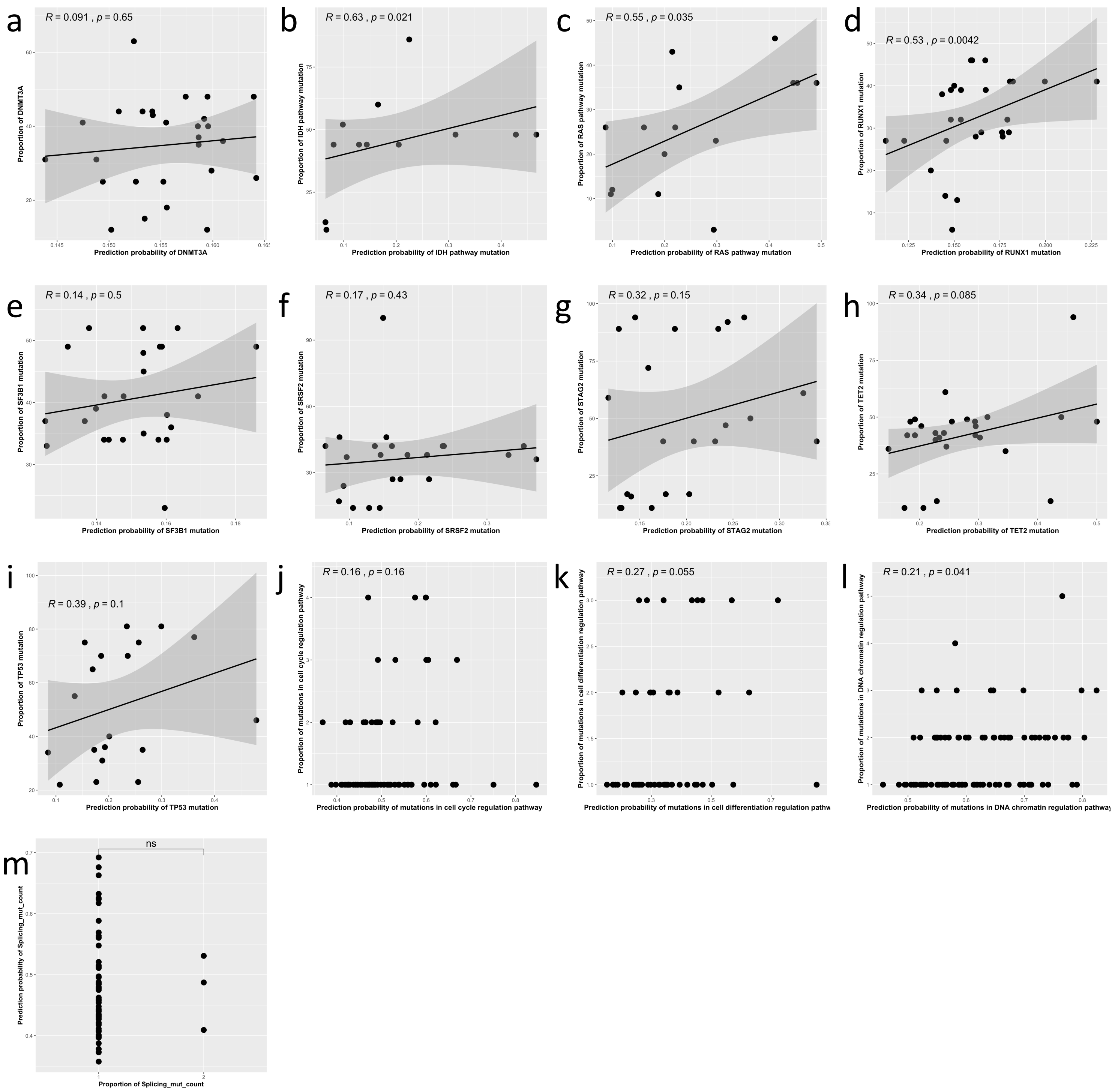


Extended Data Figure 4. Association between predicted and detected mutation occurrence. Linear regression between prediction probability of (a) *DNMT3A*, (b) *IDH1/IDH2*, (c) *KRAS/NRAS*, (d) *RUNX1*, (e) *SF3B1*, (f) *SRSF2*, (g) *STAG2*, (h) *TET2* and (i) *TP53* mutation (x-axis) and its detected variant allele frequency (y-axis). (h) Linear regression for predicted (j) cell cycle, (k) cell differentiation and (l) DNA chromatin structure regulation pathway mutation probability and detected frequency of altered genes in respective pathway. The R correlation coefficient value has been computed with Spearman correlation. (m) Wilcoxon comparison for predicted spliceosome regulation pathway mutation probability and detected frequency of altered genes in the *RAS* pathway.

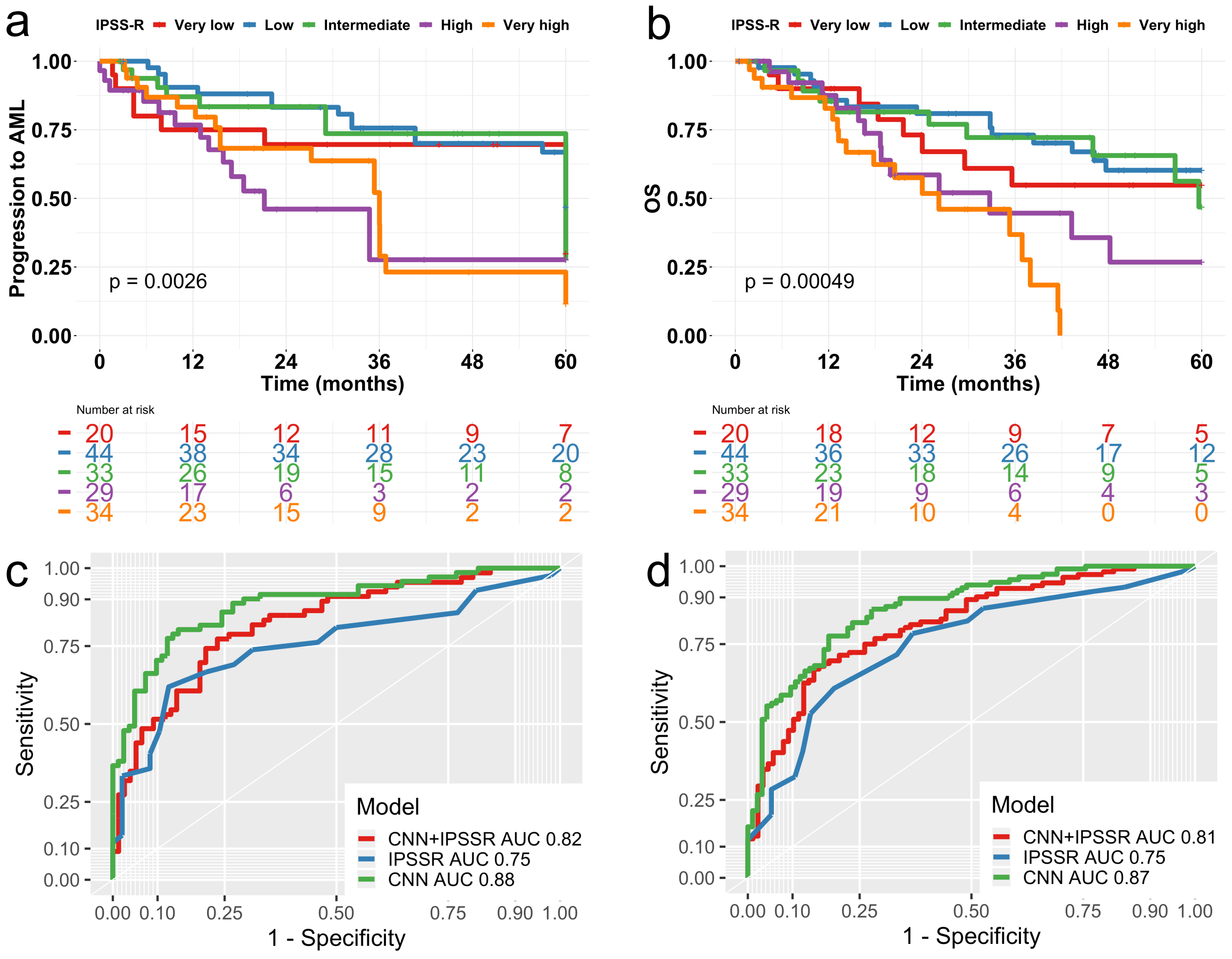


Extended Data Figure 5. (a) Progression to AML and (b) overall survival stratified with IPSSR. Kaplan-Meier curves are compared with Cox regression analysis (log-rank test). (c-d) Receiver operating characteristic (ROC) curves for models developed with features extracted from convolutional neural networks (CNN), Revised International Prognostic Scoring System (IPSSR) or their combination (CNN+IPSSR) to predict (c) progression to AML in 2 years or (d) overall survival in 2 years.

Supplementary Figure 6. Model reports for prediction of (a) MDS vs. MDS/MPN, (b) *ASXL1* mutation, (c) chromosome 5q deletion, (d) abnormal karyotype, (e) progression to AML in 2 years, (f) mutation in genes regulating cell cycle, (g) mutation in genes regulating cell differentiation, (h) complex karyotype, (i) chromosome 7q deletion, (j) chromosome 20q deletion, (k) mutation in genes regulating DNA chromatin structure, (l) *DNMT3A* mutation, (m) female gender, (n) *IDH1/IDH2* mutation, (o) chromosome 7q monosomy, (p) overall survival in 2 years, (q) *KRAS/NRAS* mutation, (r) mutation in genes regulating *RAS* pathway, (s) *RUNX1* mutation, (t) secondary MDS, (u) *SF3B1* mutation, (v) mutation in genes regulating splicesome, (w) *SRSF2* mutation, (x) *STAG2* mutation, (y) *TET2* mutation, (z) *TP53* mutation, and (aa) chromosome 8 trisomy.

Reports include plots for (i) Variable numbers of the elastic net regression models with least crossvalidation error and best-performing penalization level. (ii) The plot with highest area under the receiver operating characteristic (AUROC) curve to evaluate tile-level label and (iii) TMA spot level label prediction. (iv) Two representative tile-level and TMA spot level images and activation maps visualizing the prediction probability.

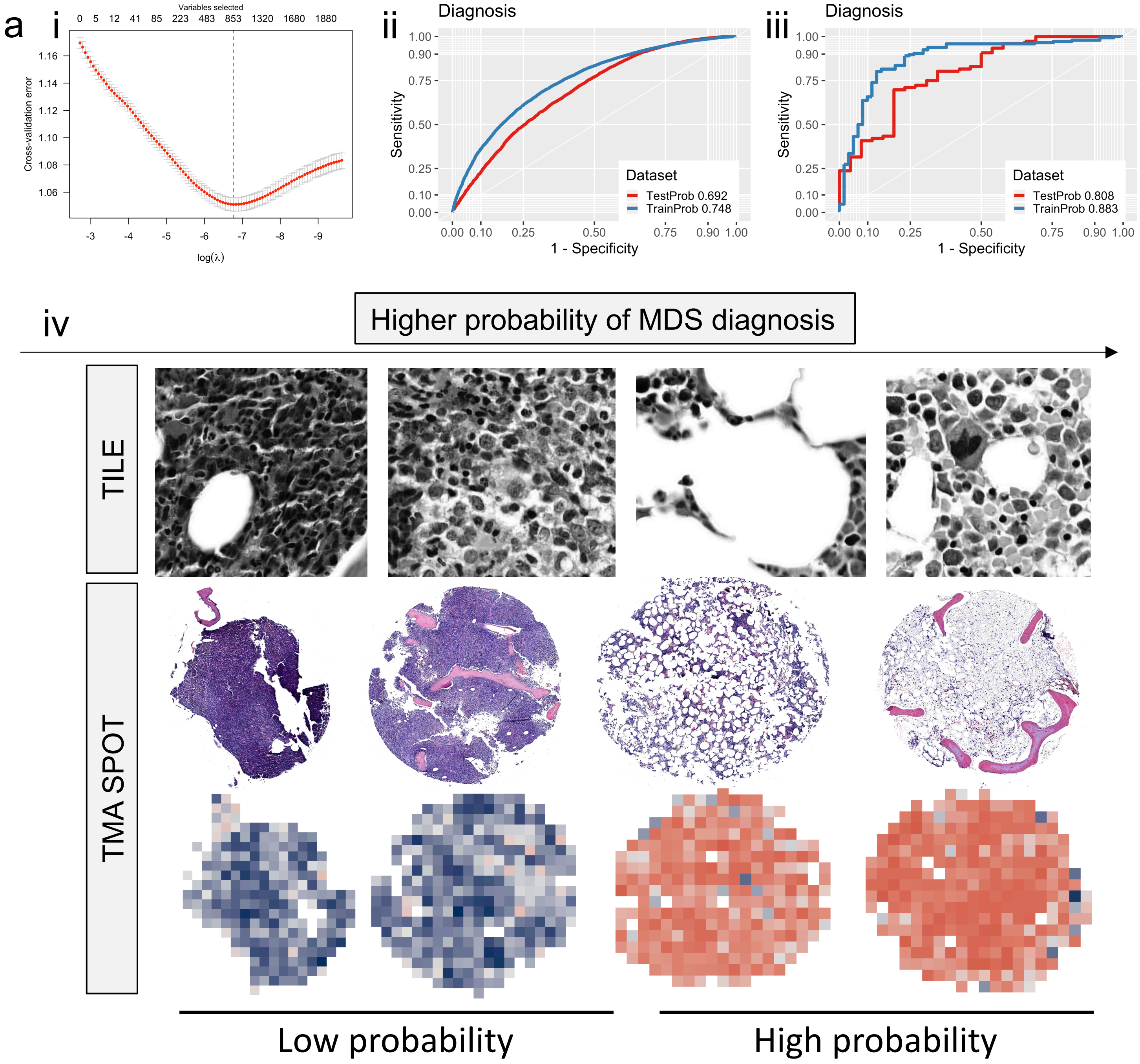


Extended Data Figure 6a.

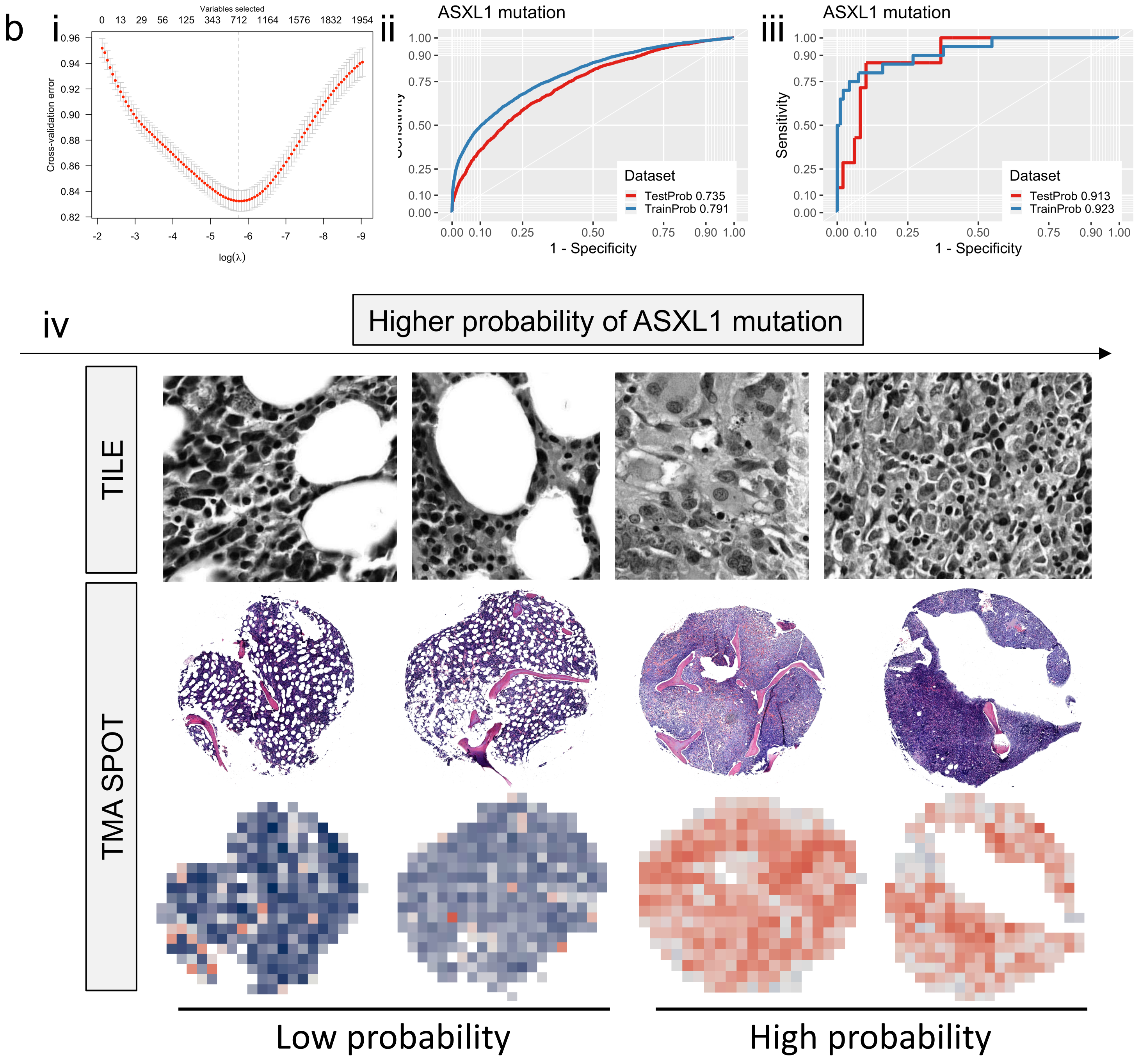


Extended Data Figure 6b.

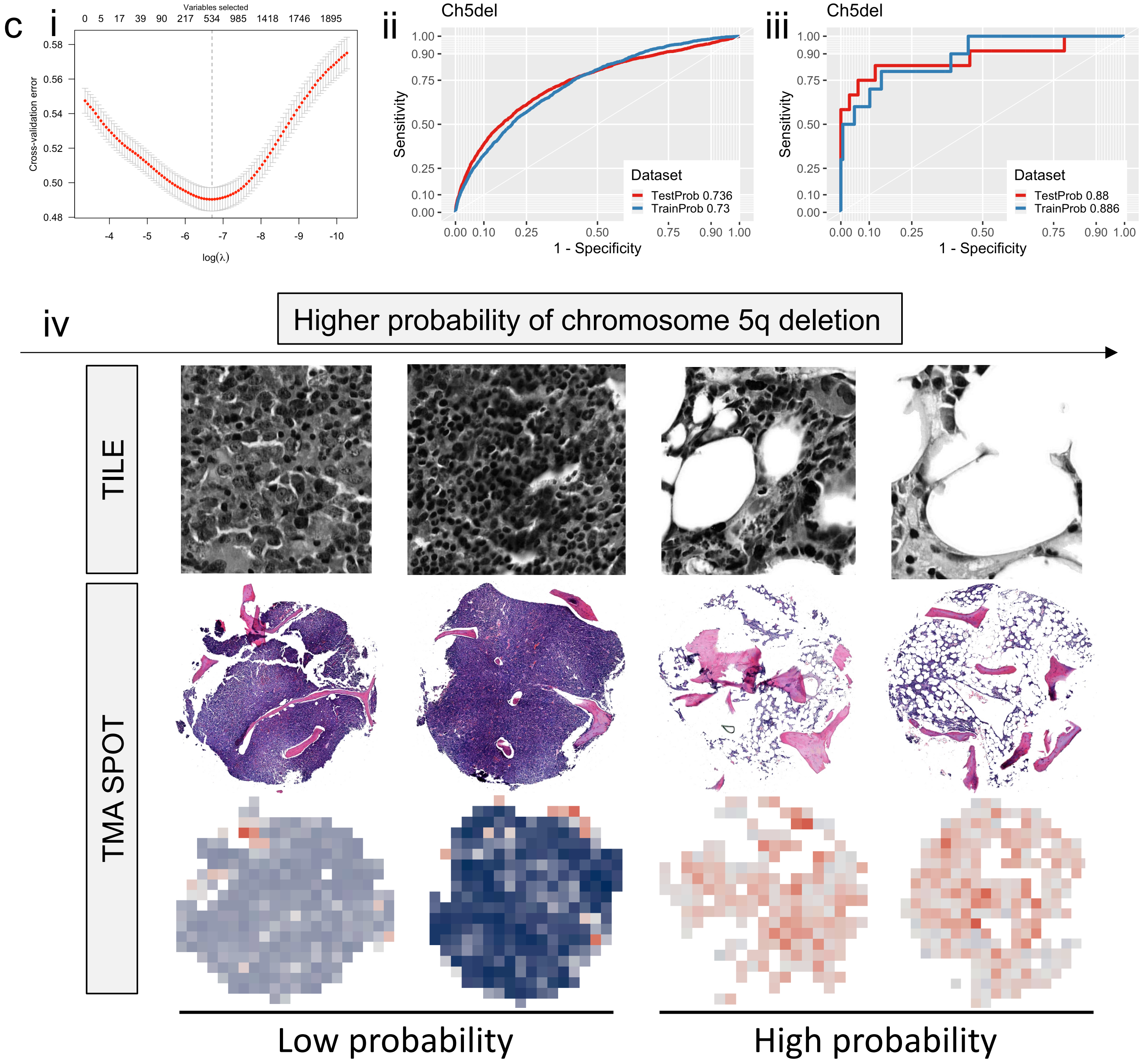


Extended Data Figure 6c.

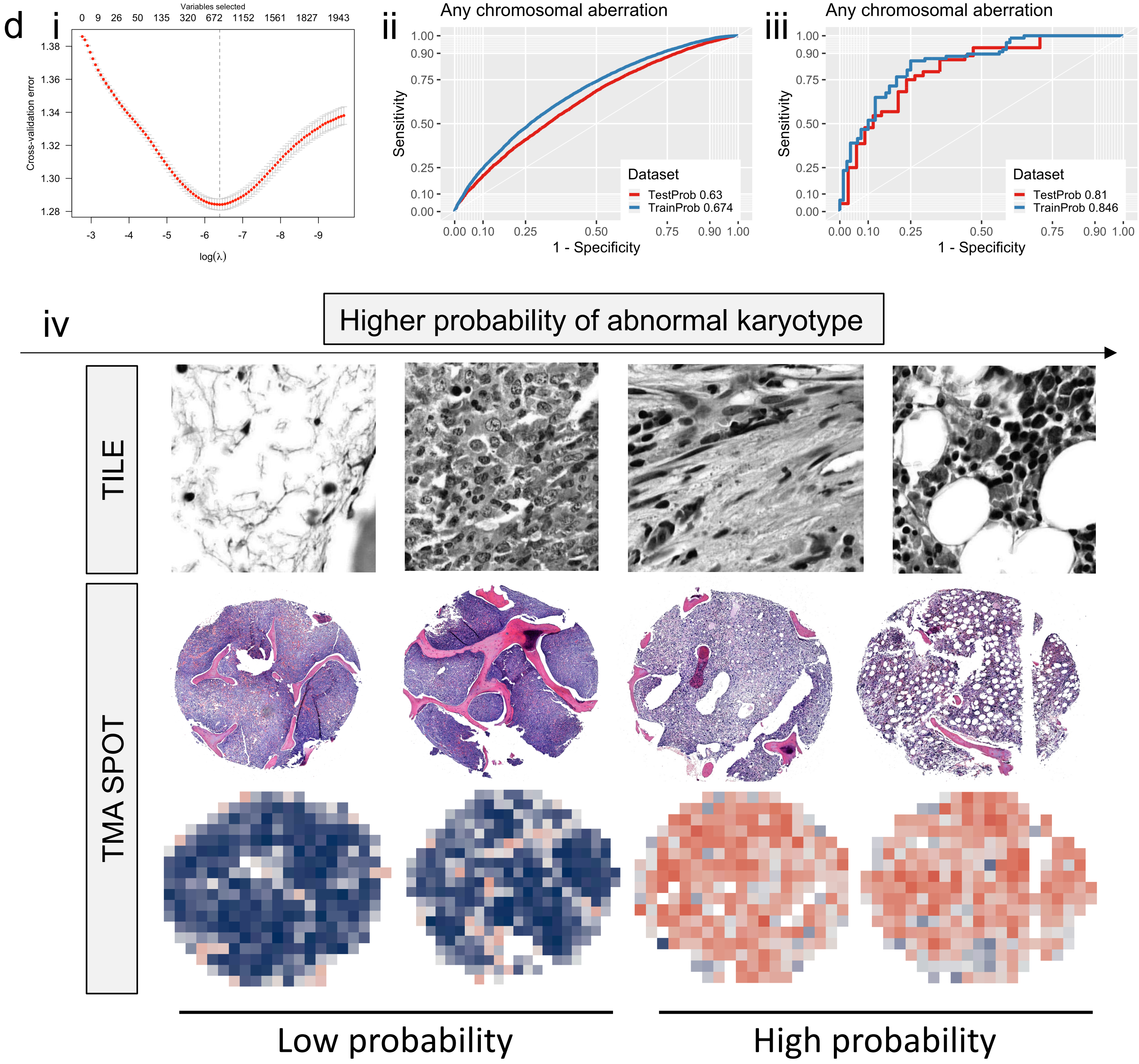


Extended Data Figure 6d.

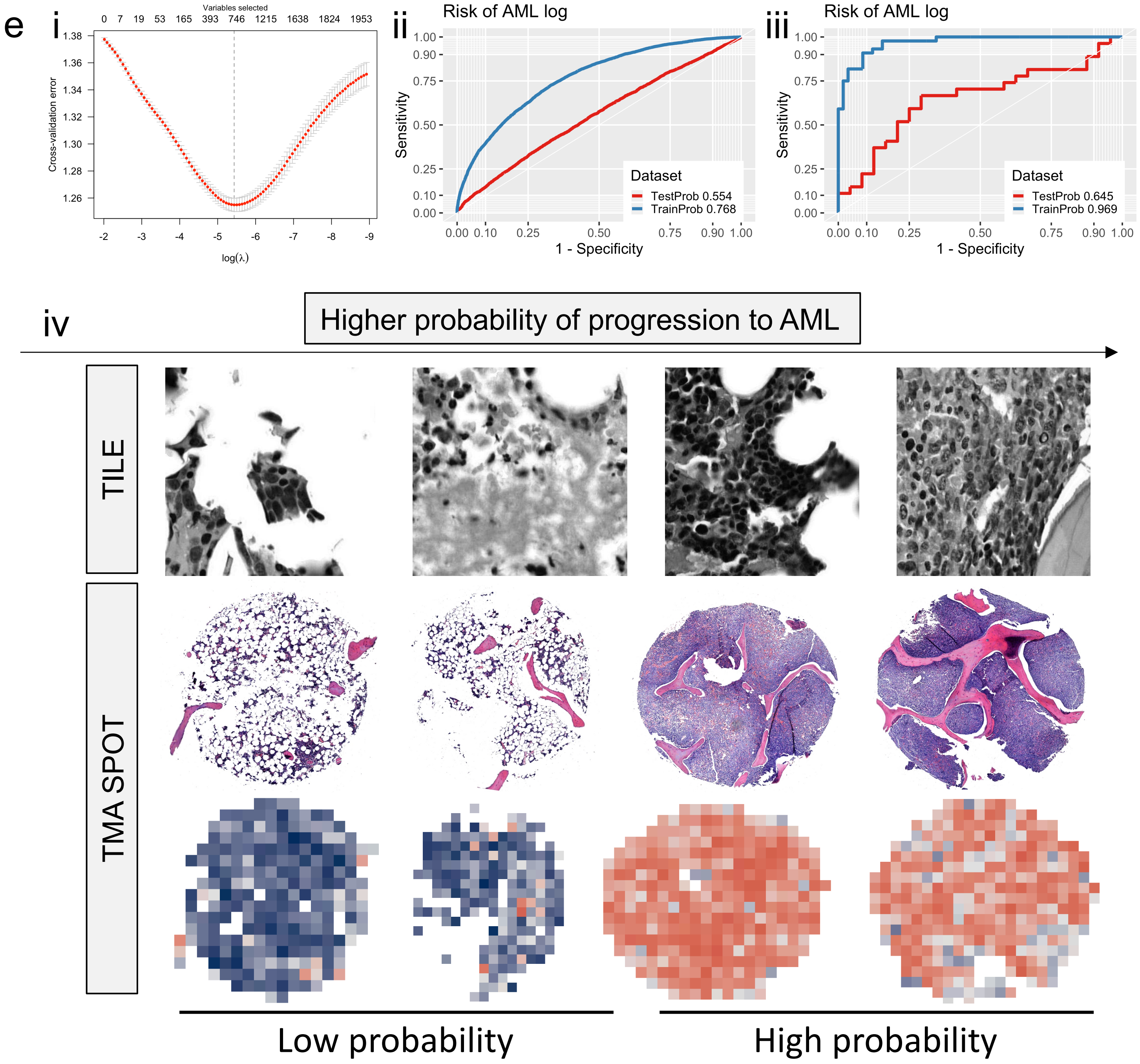


Extended Data Figure 6e.

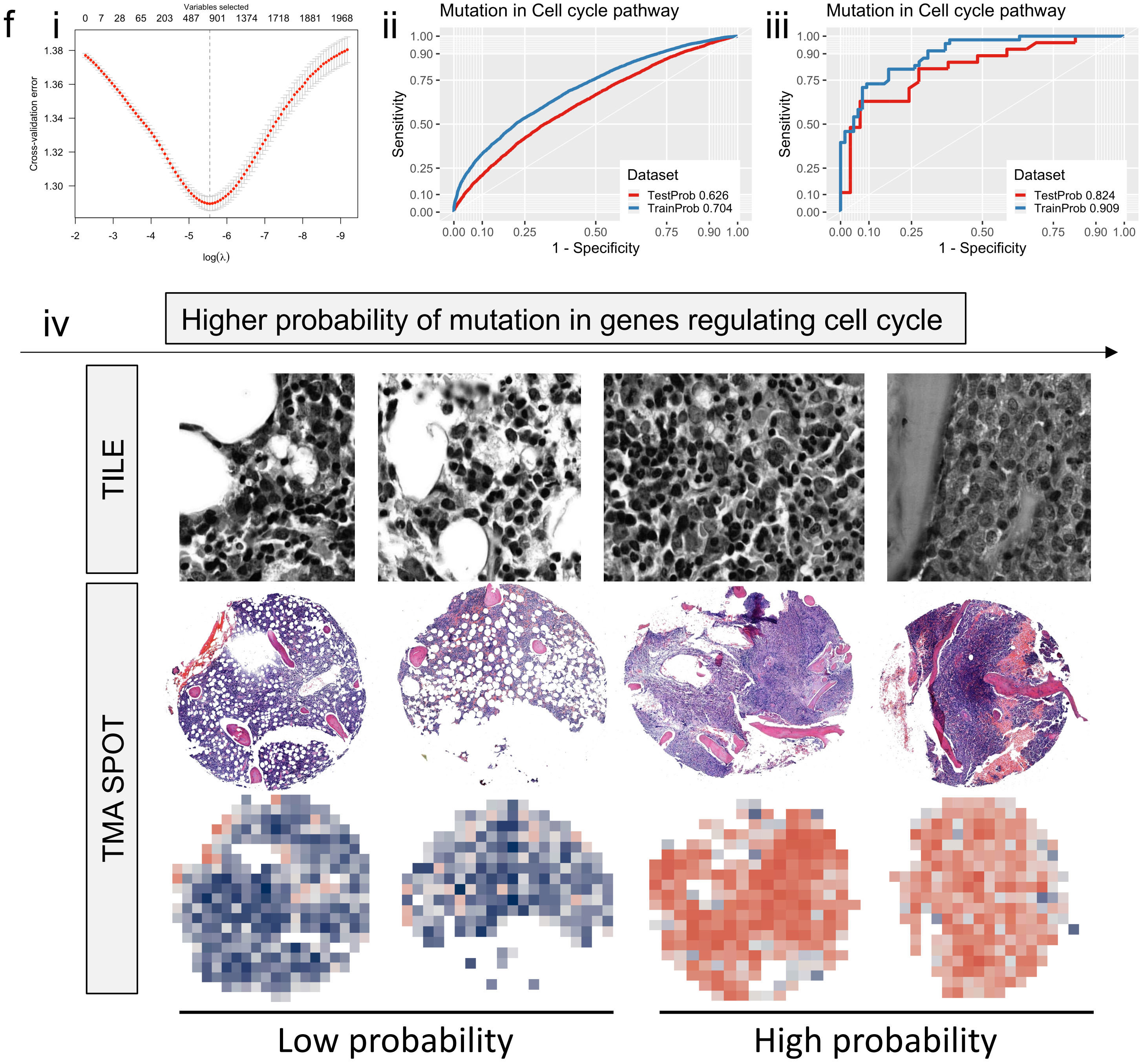


Extended Data Figure 6f.

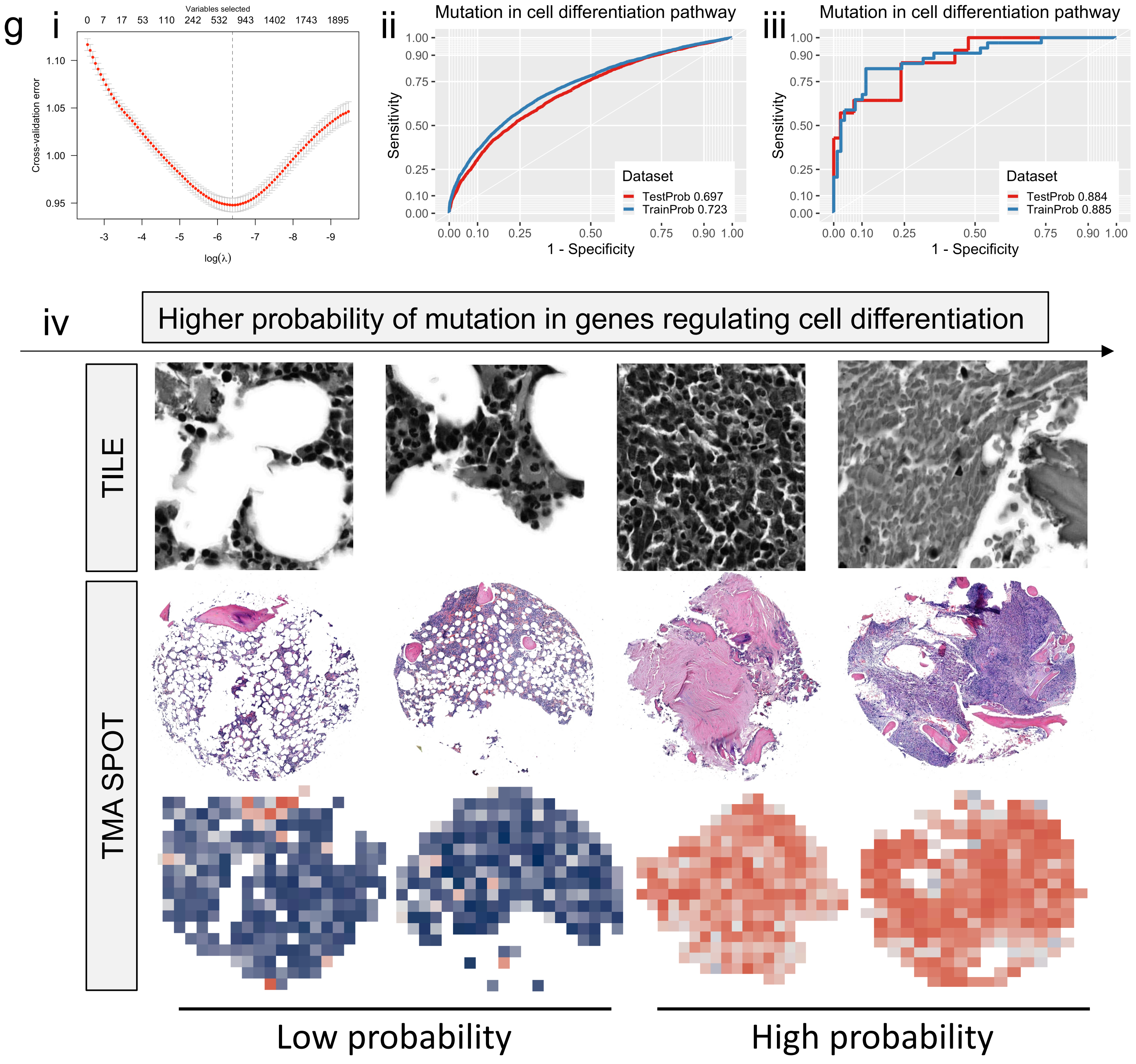


Extended Data Figure 6g.

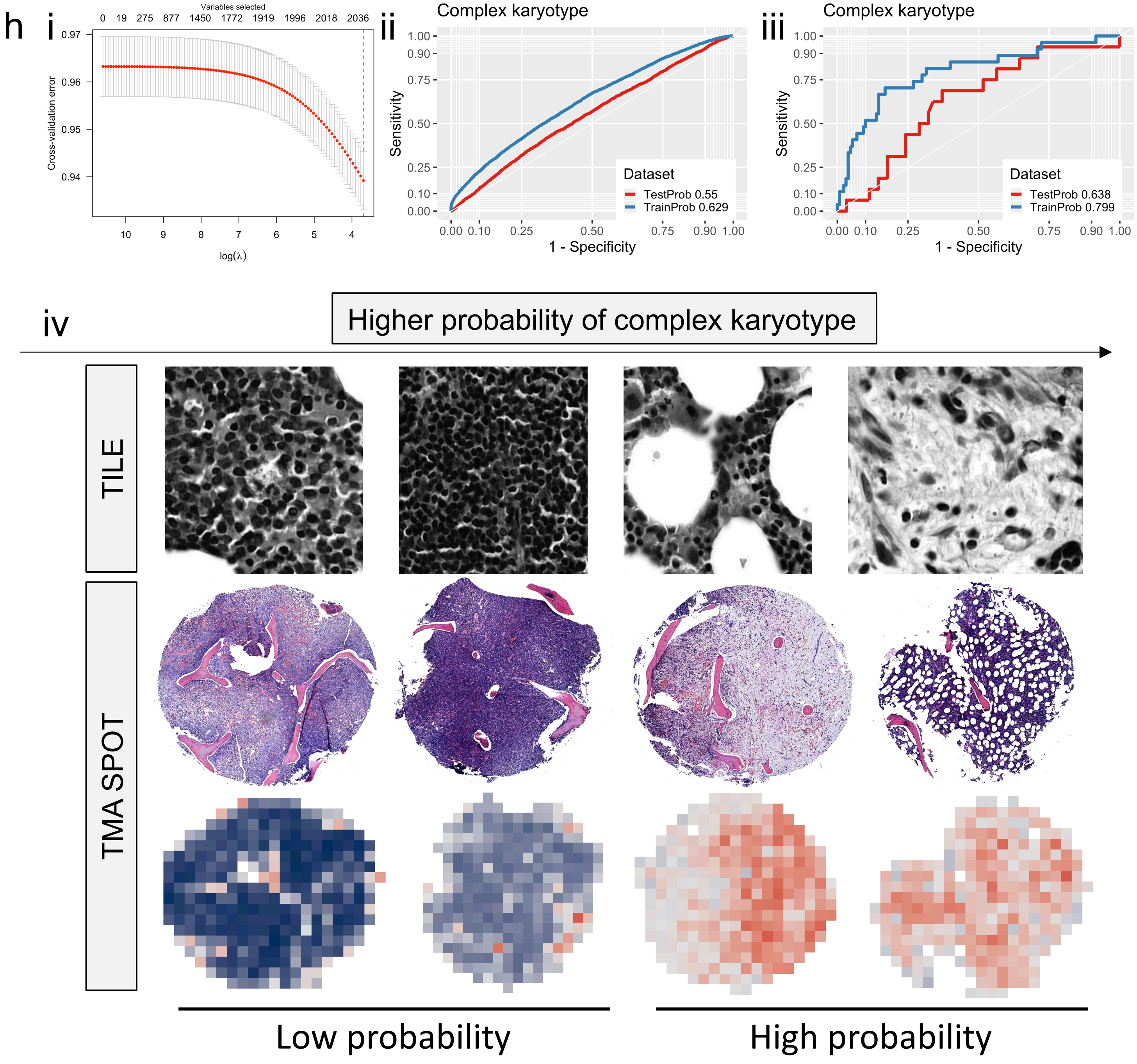


Extended Data Figure 6h.

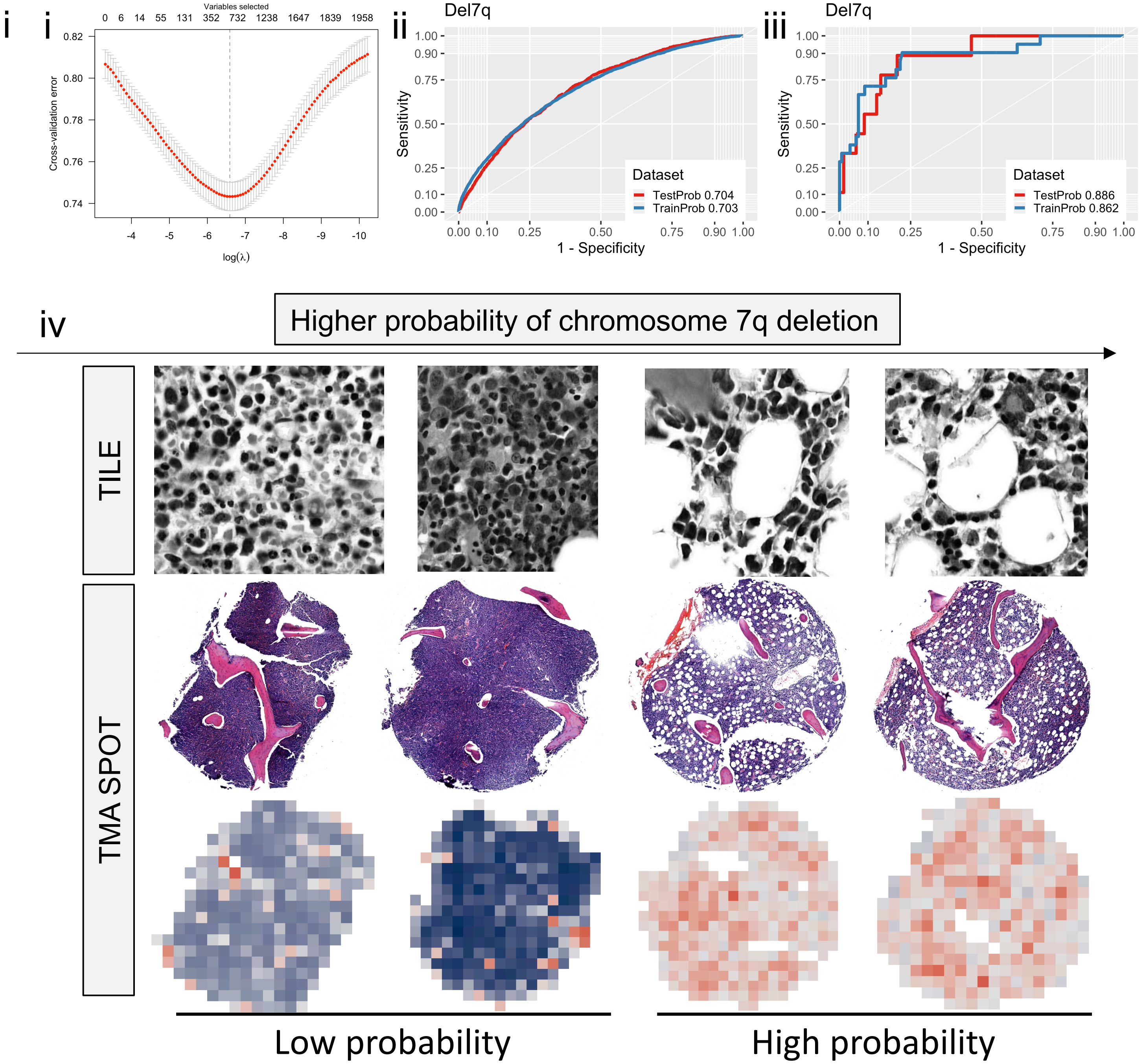


Extended Data Figure 6i.

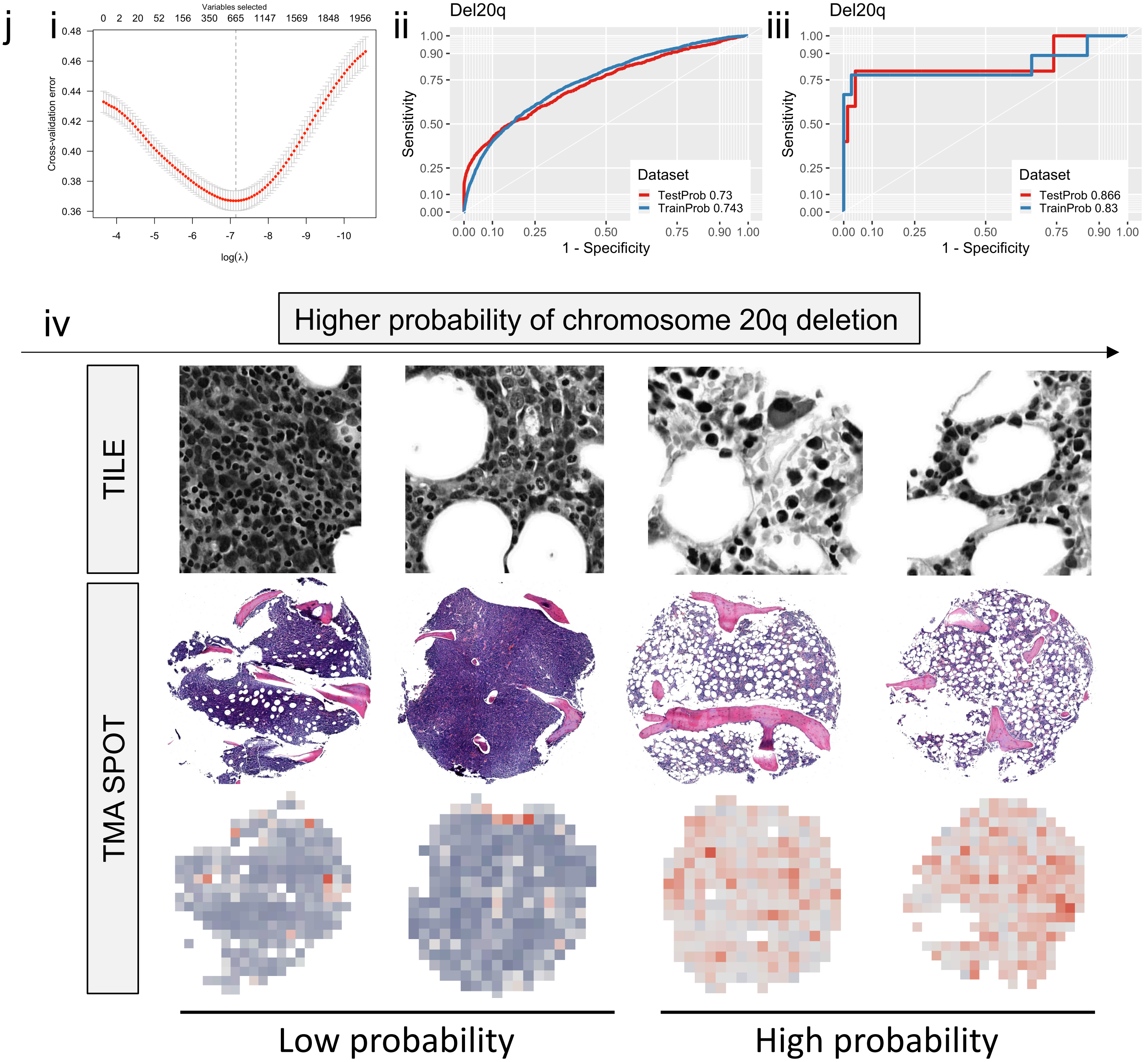


Extended Data Figure 6j.

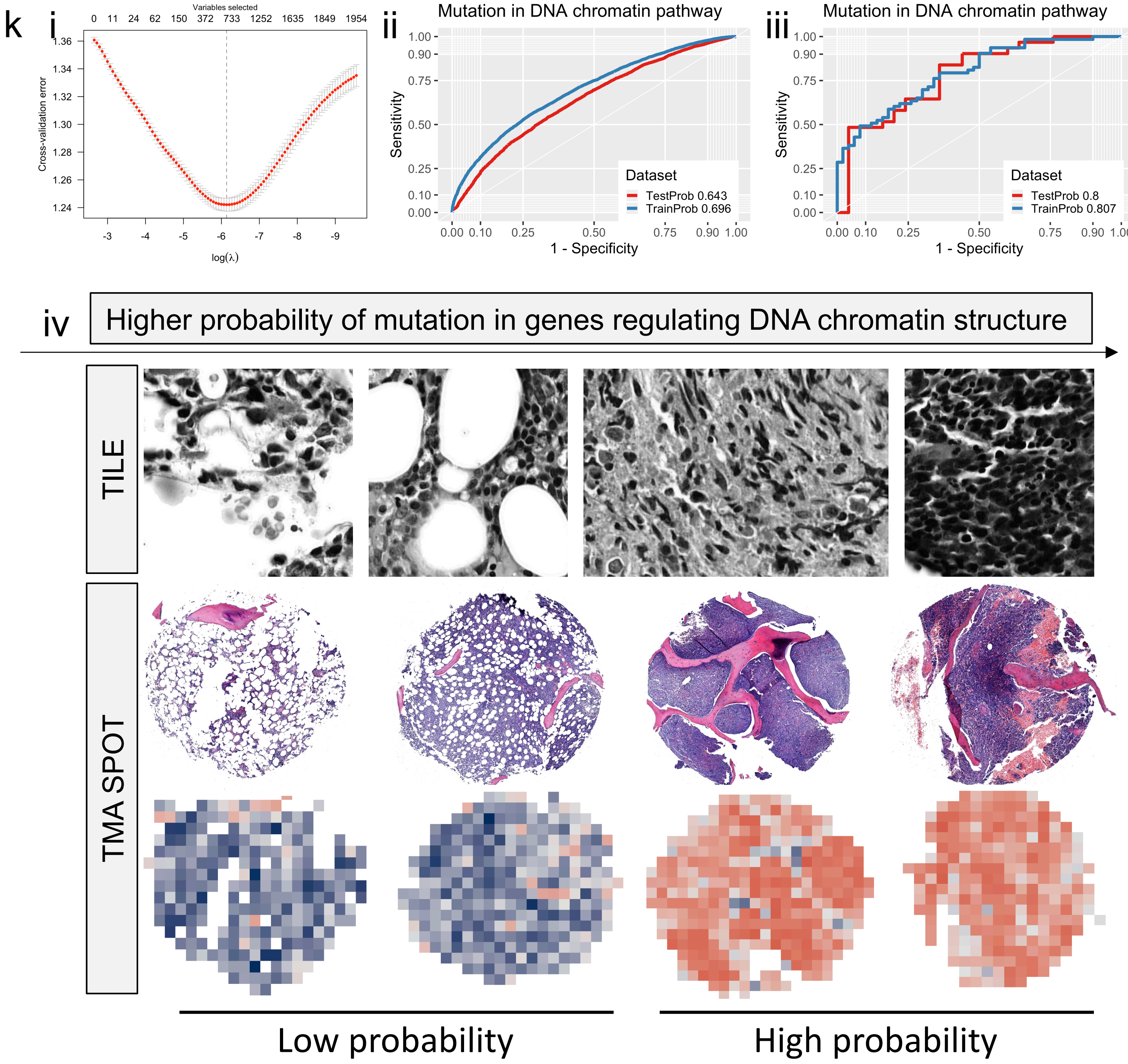


Extended Data Figure 6k.

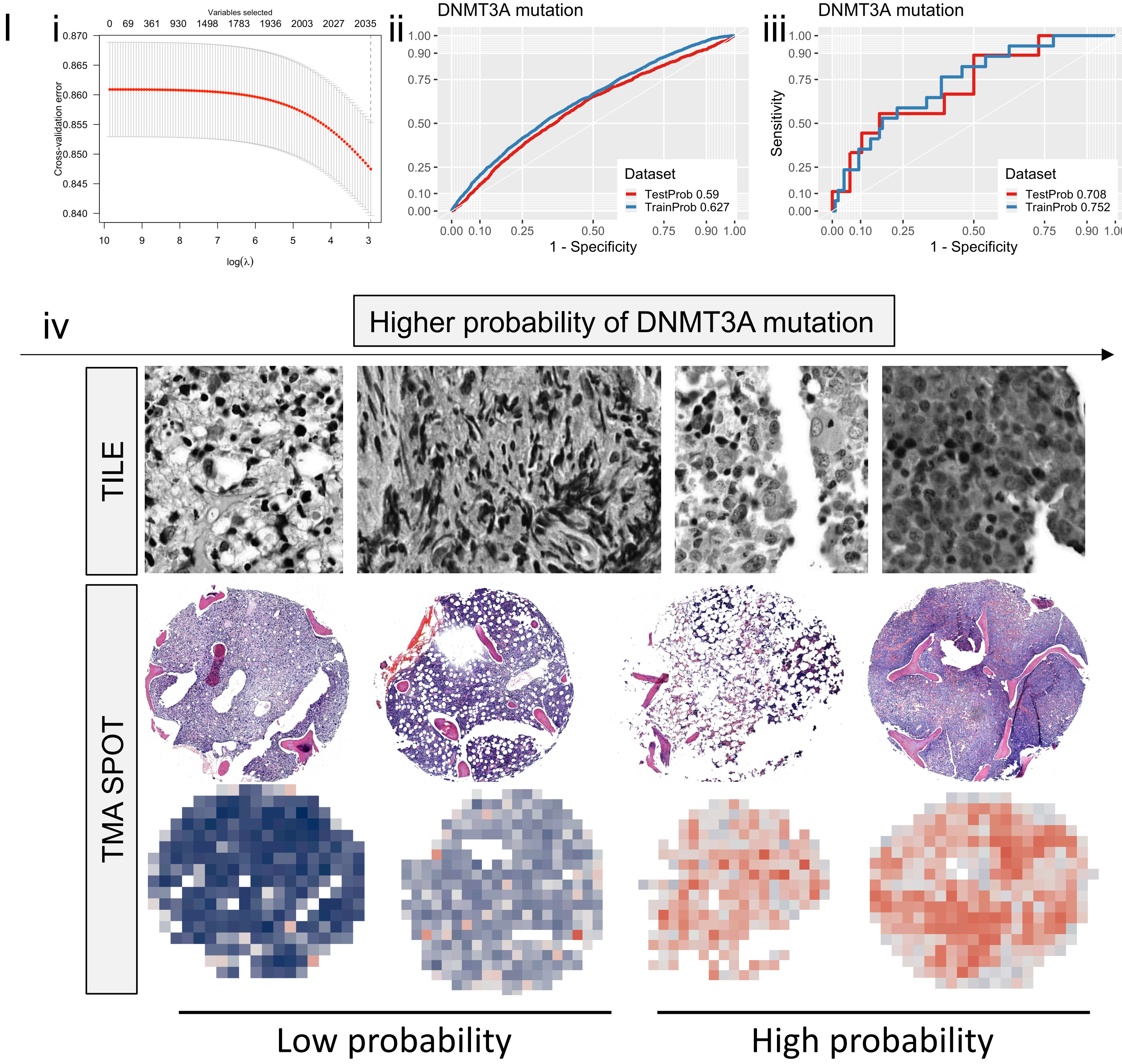


Extended Data Figure 6l.

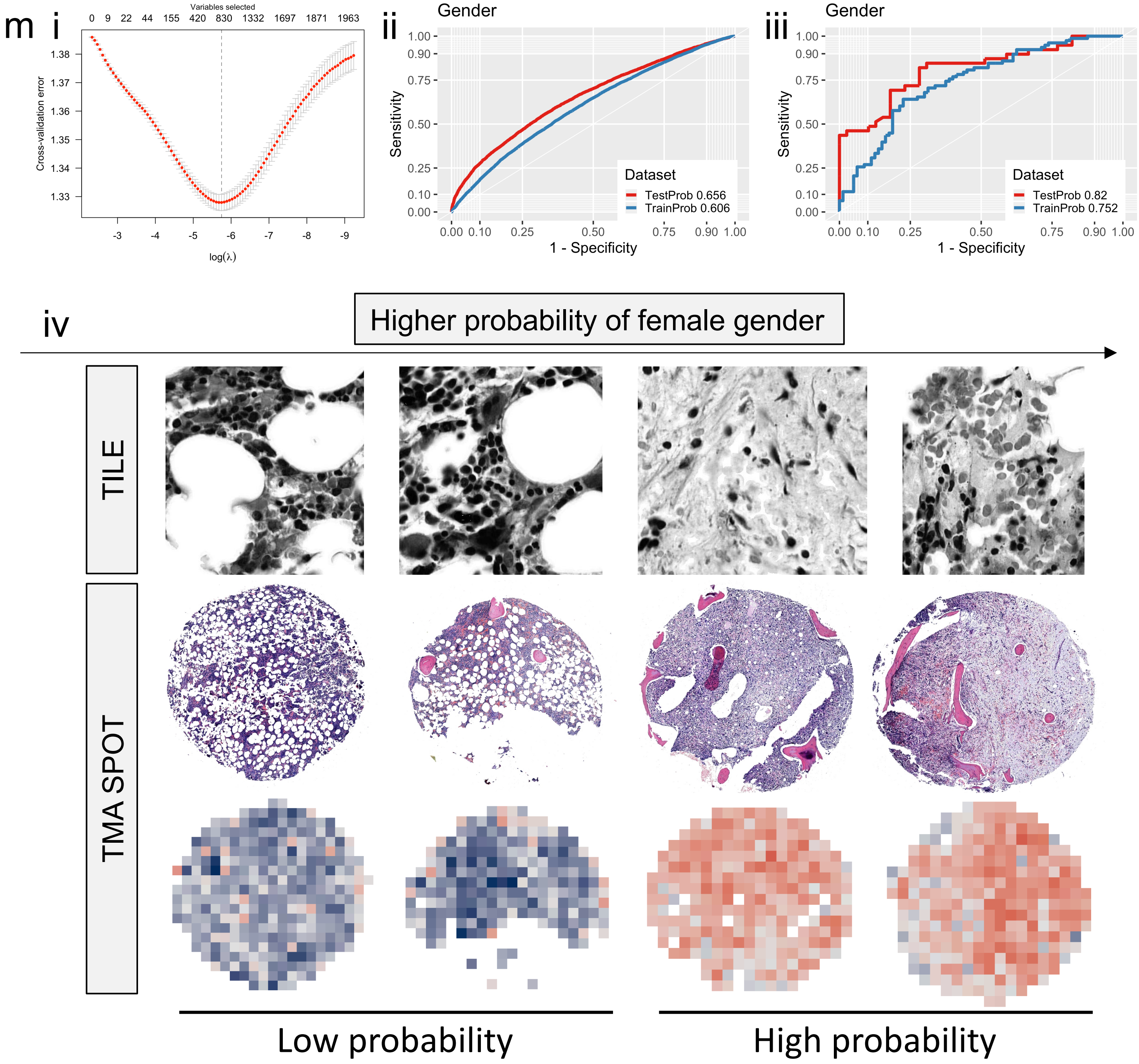


Extended Data Figure 6m.

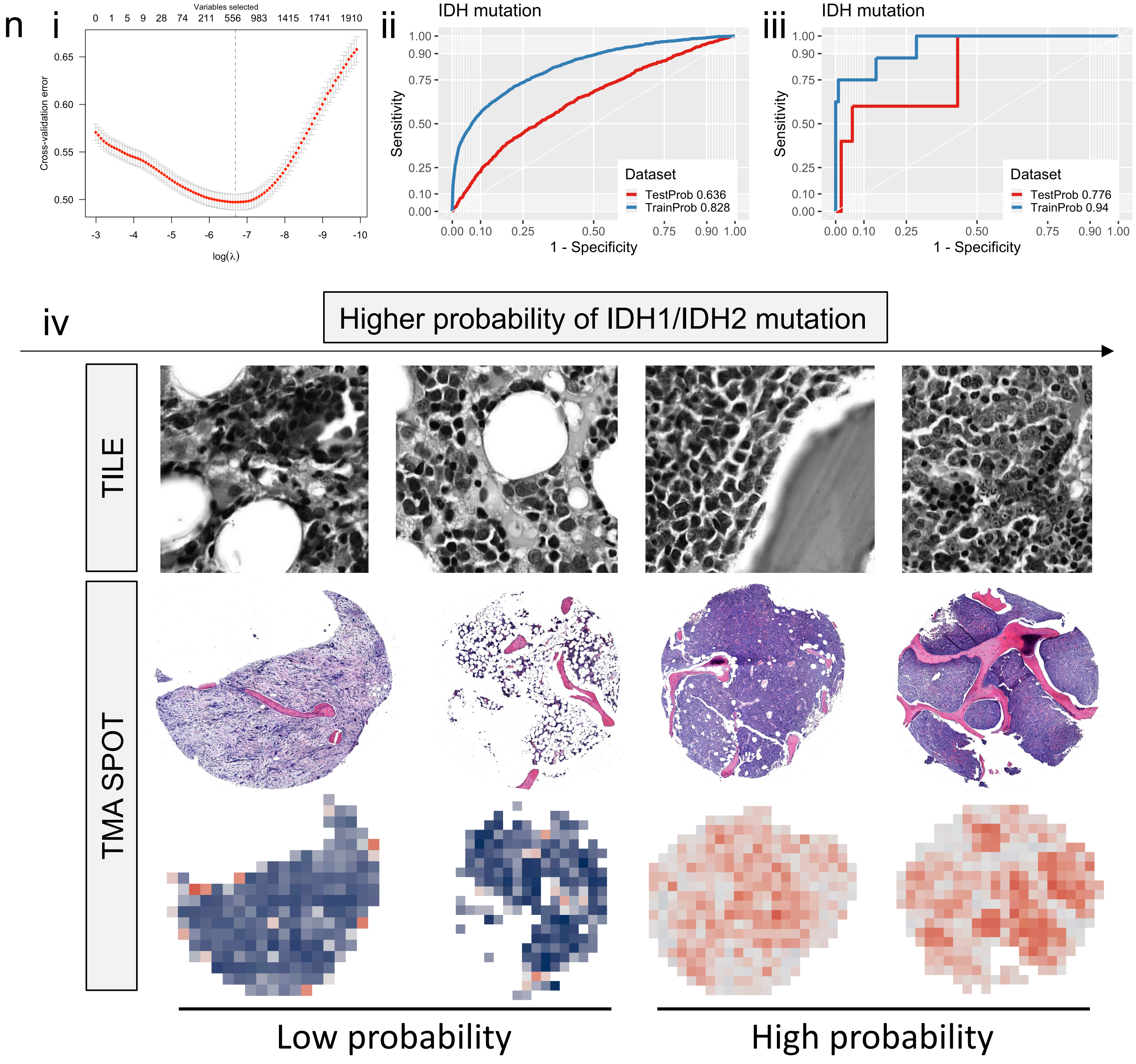


Extended Data Figure 6n.

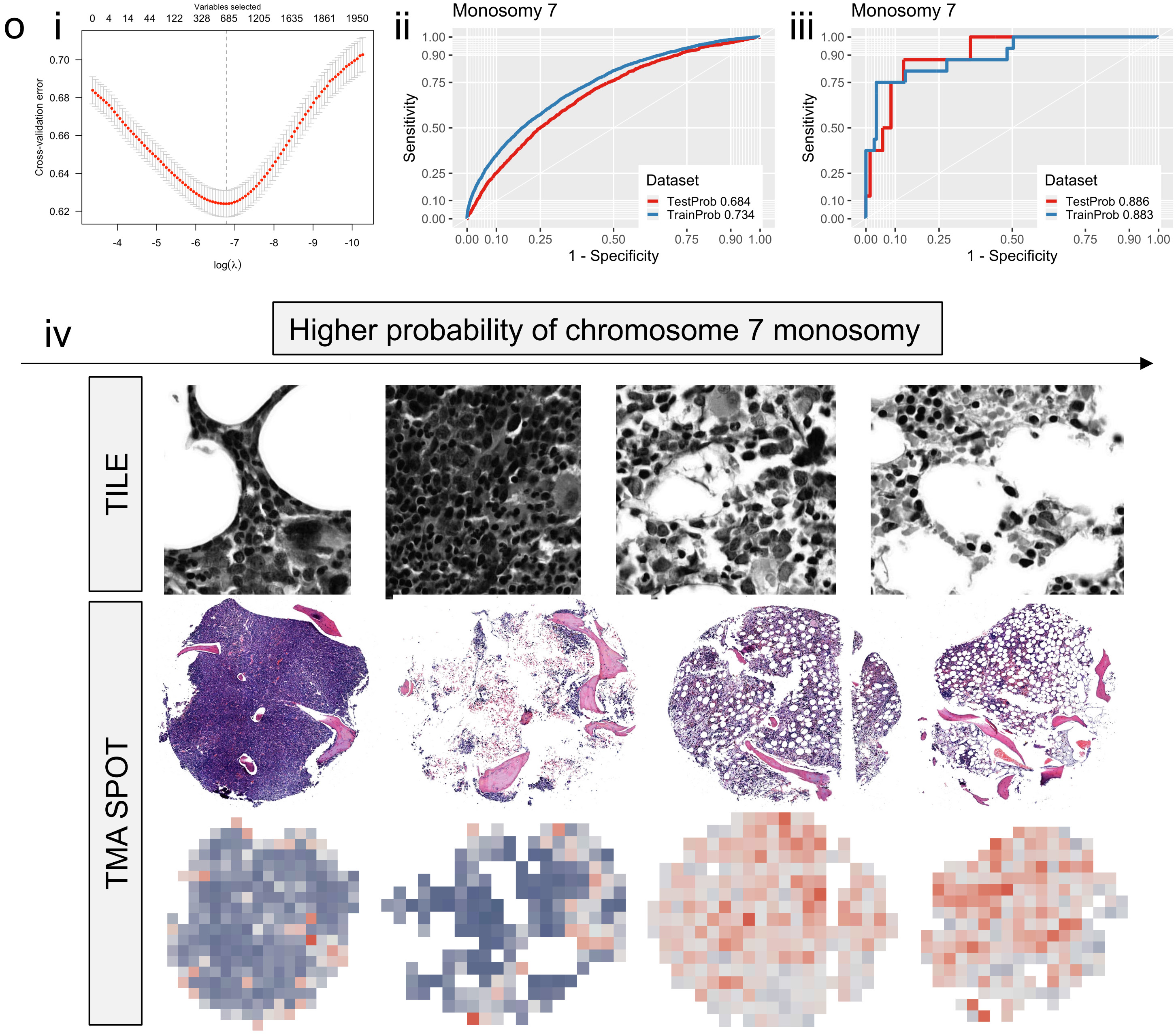


Extended Data Figure 6o.

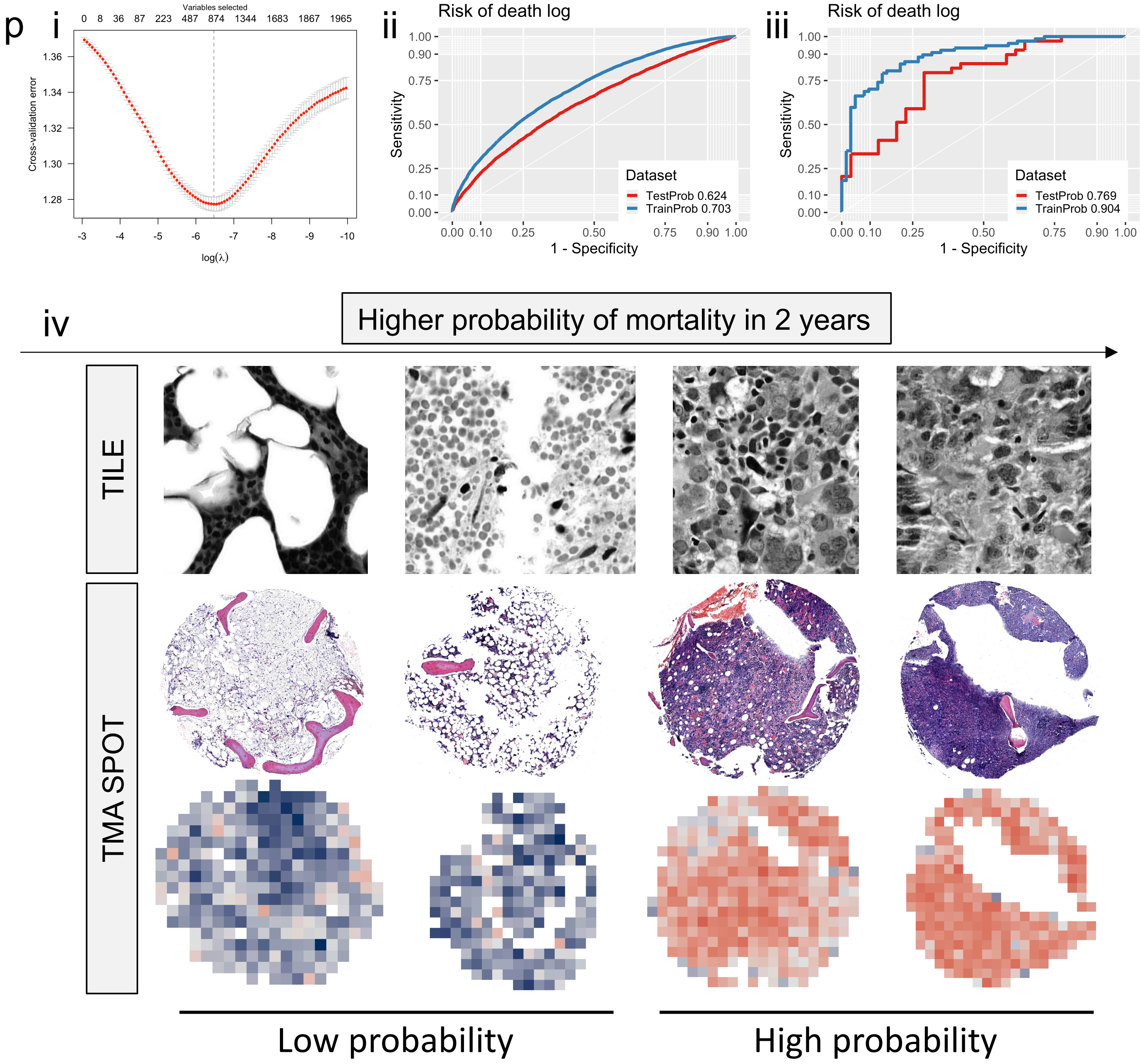


Extended Data Figure 6p.

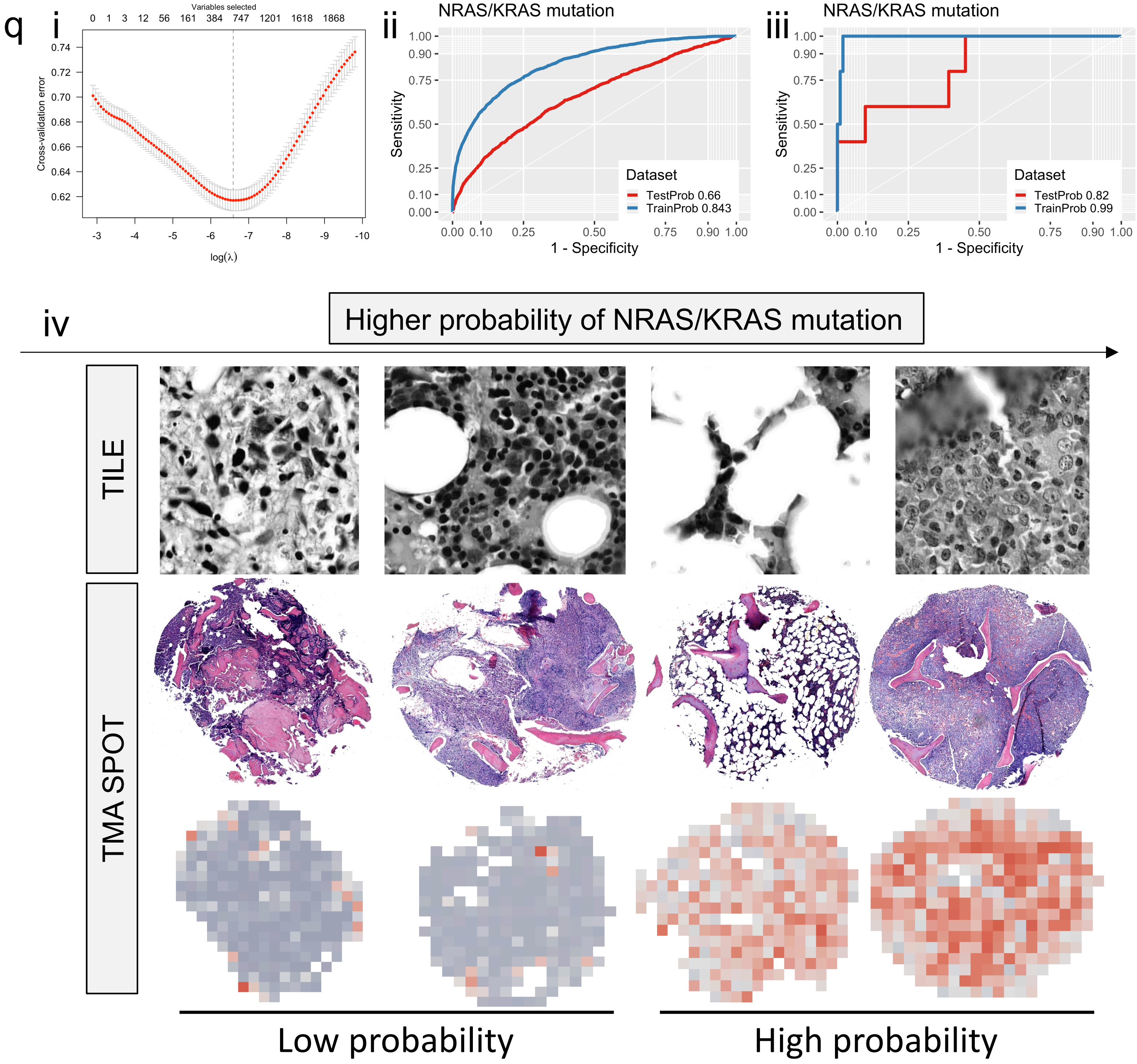


Extended Data Figure 6q.

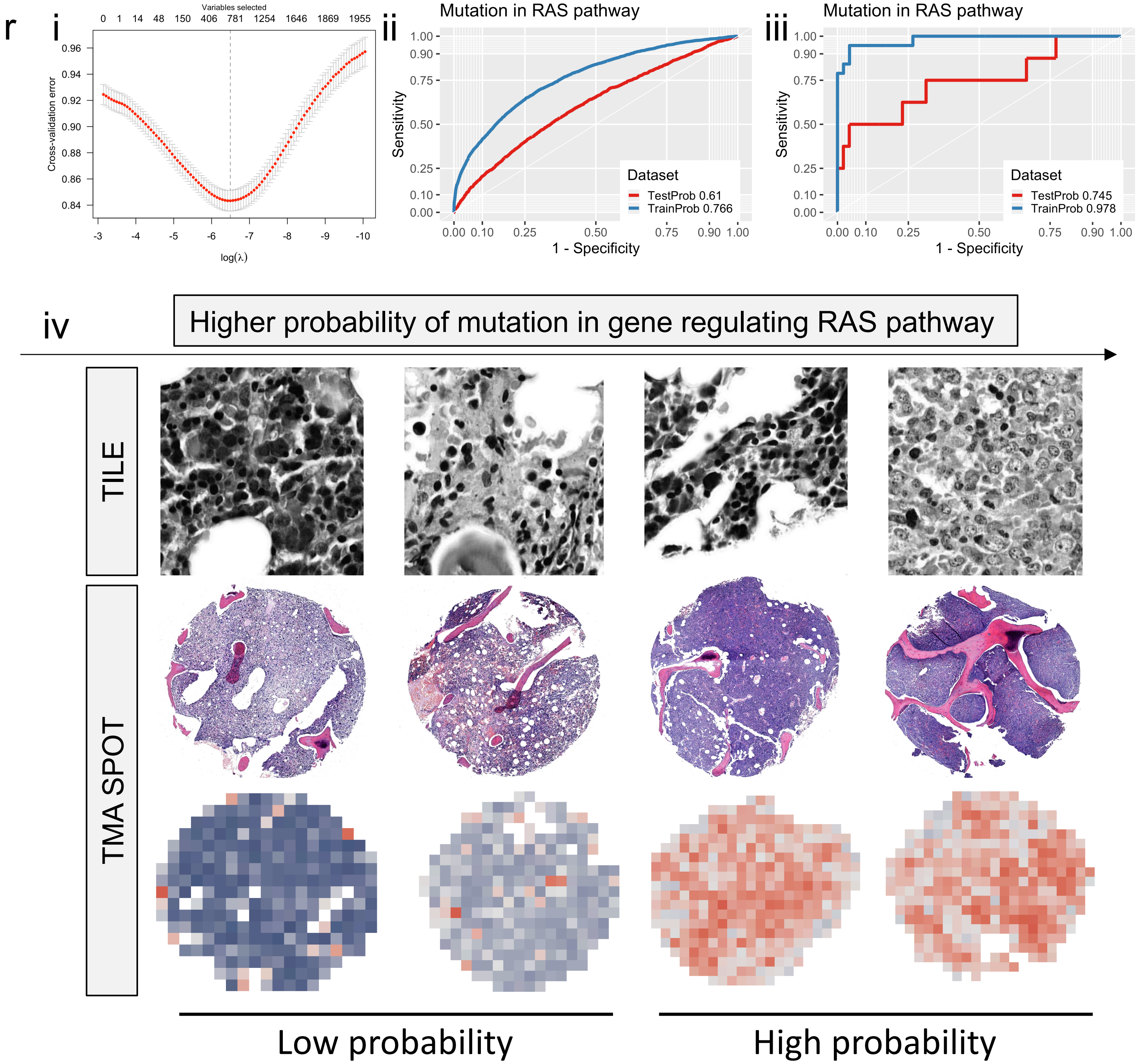


Extended Data Figure 6r.

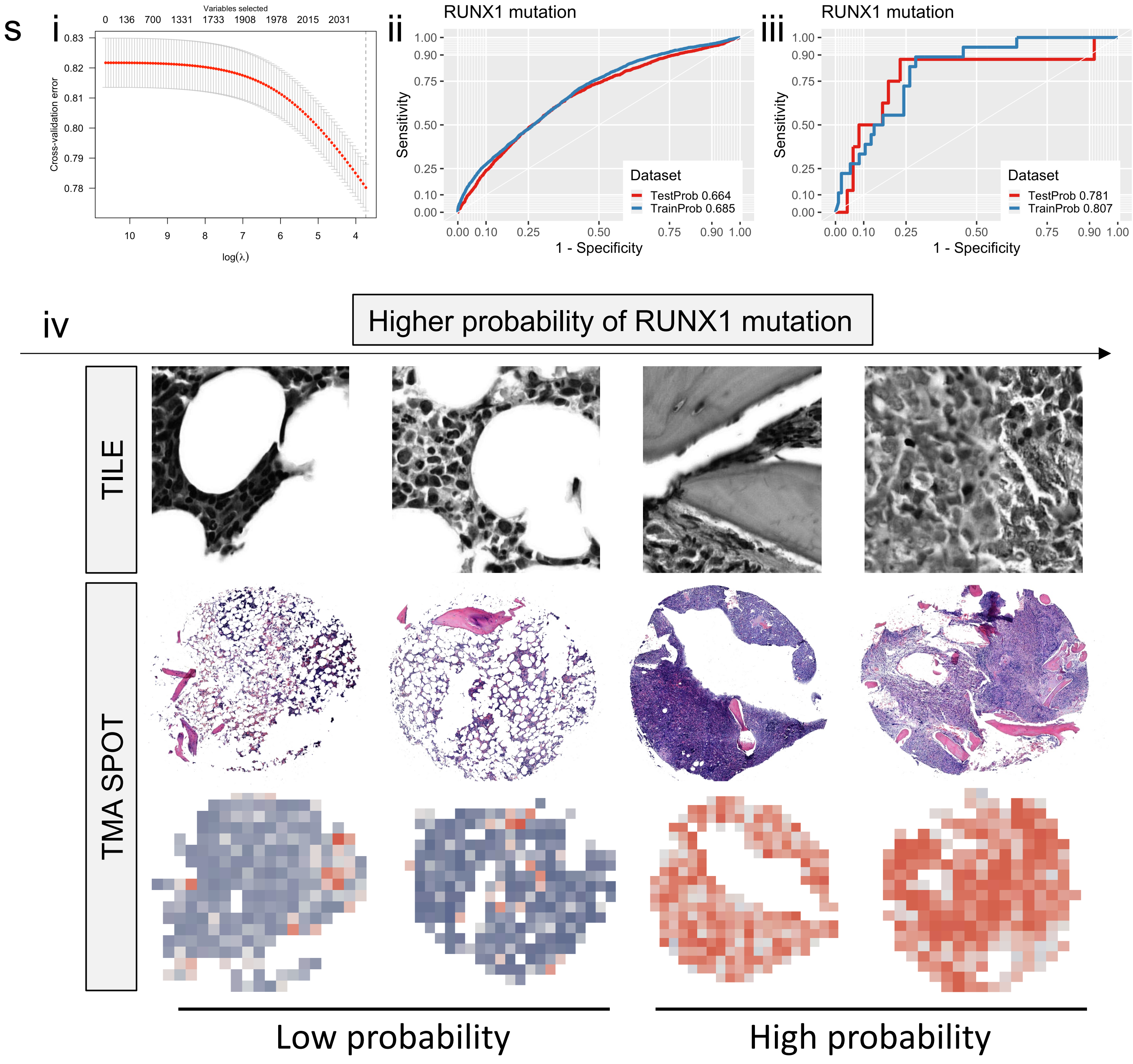


Extended Data Figure 6s.

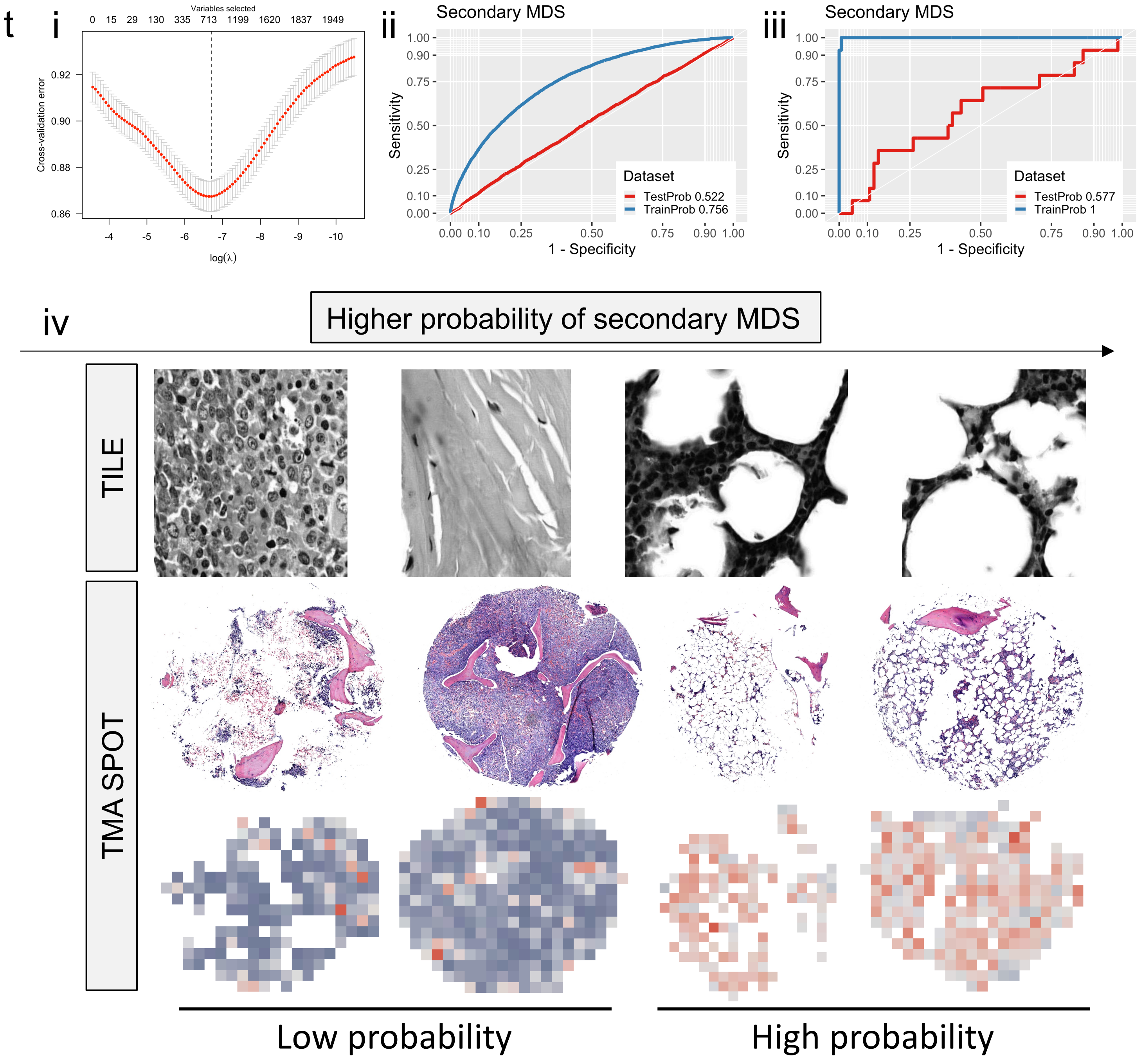


Extended Data Figure 6t.

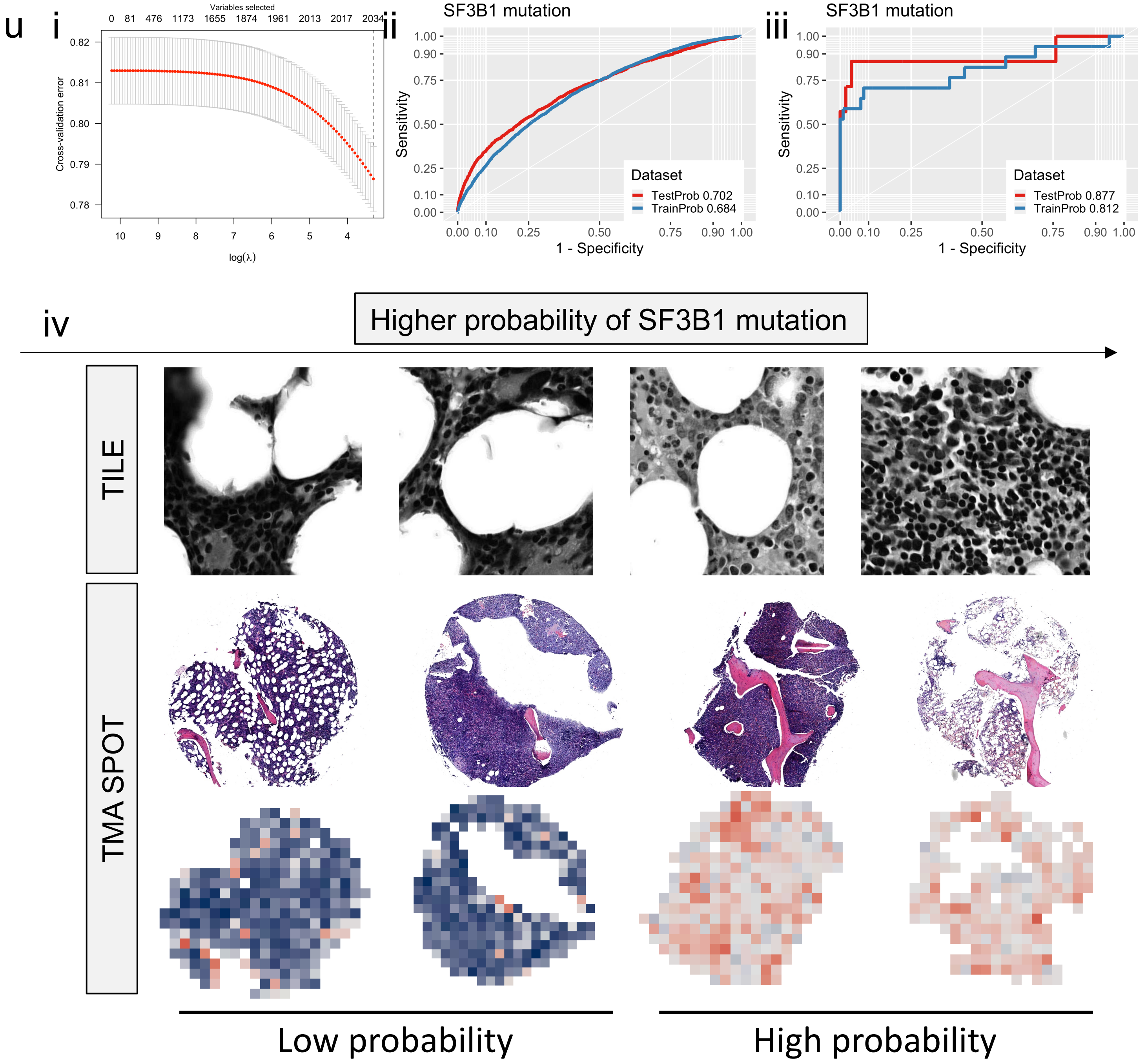


Extended Data Figure 6u.

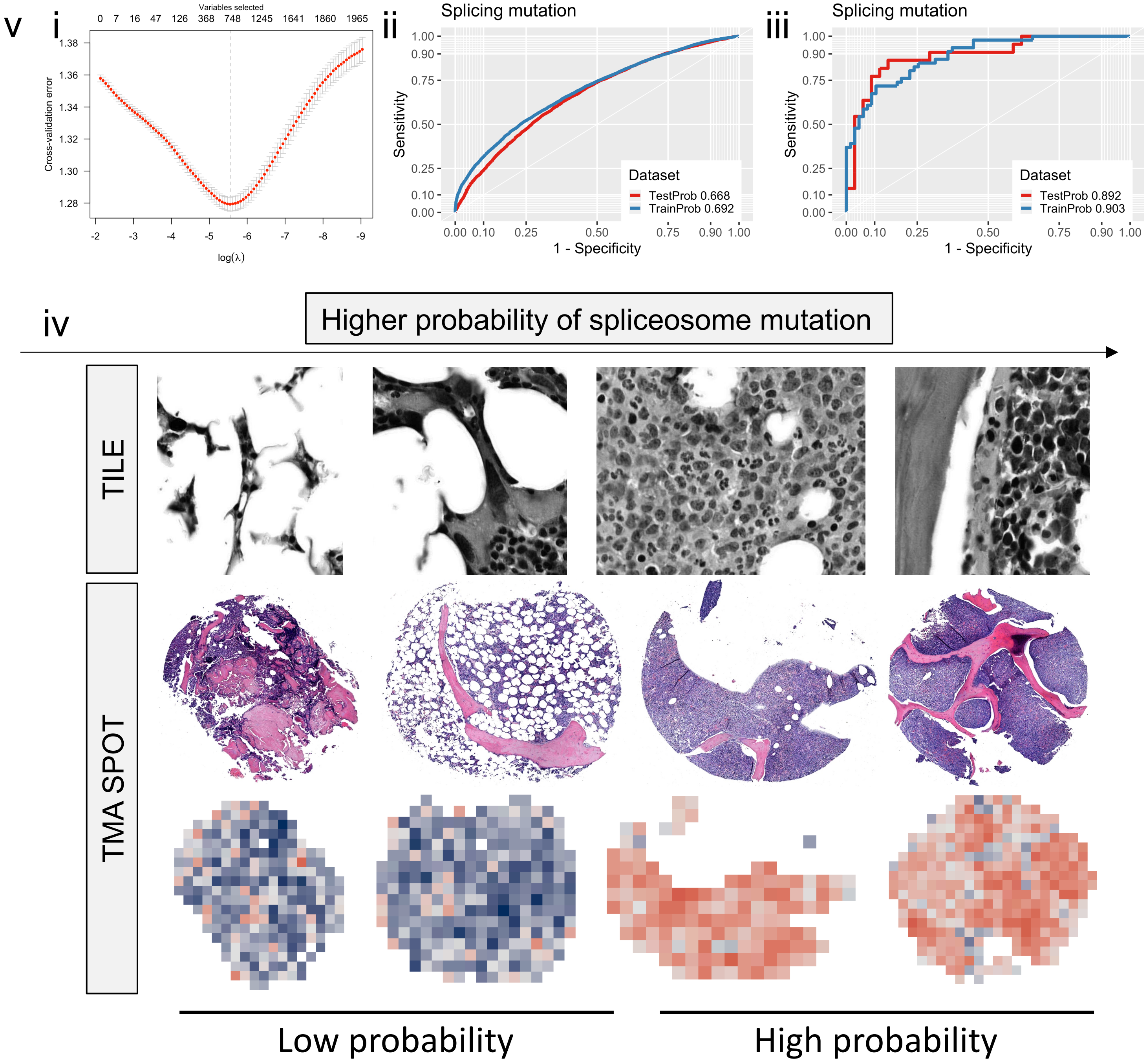


Extended Data Figure 6v.

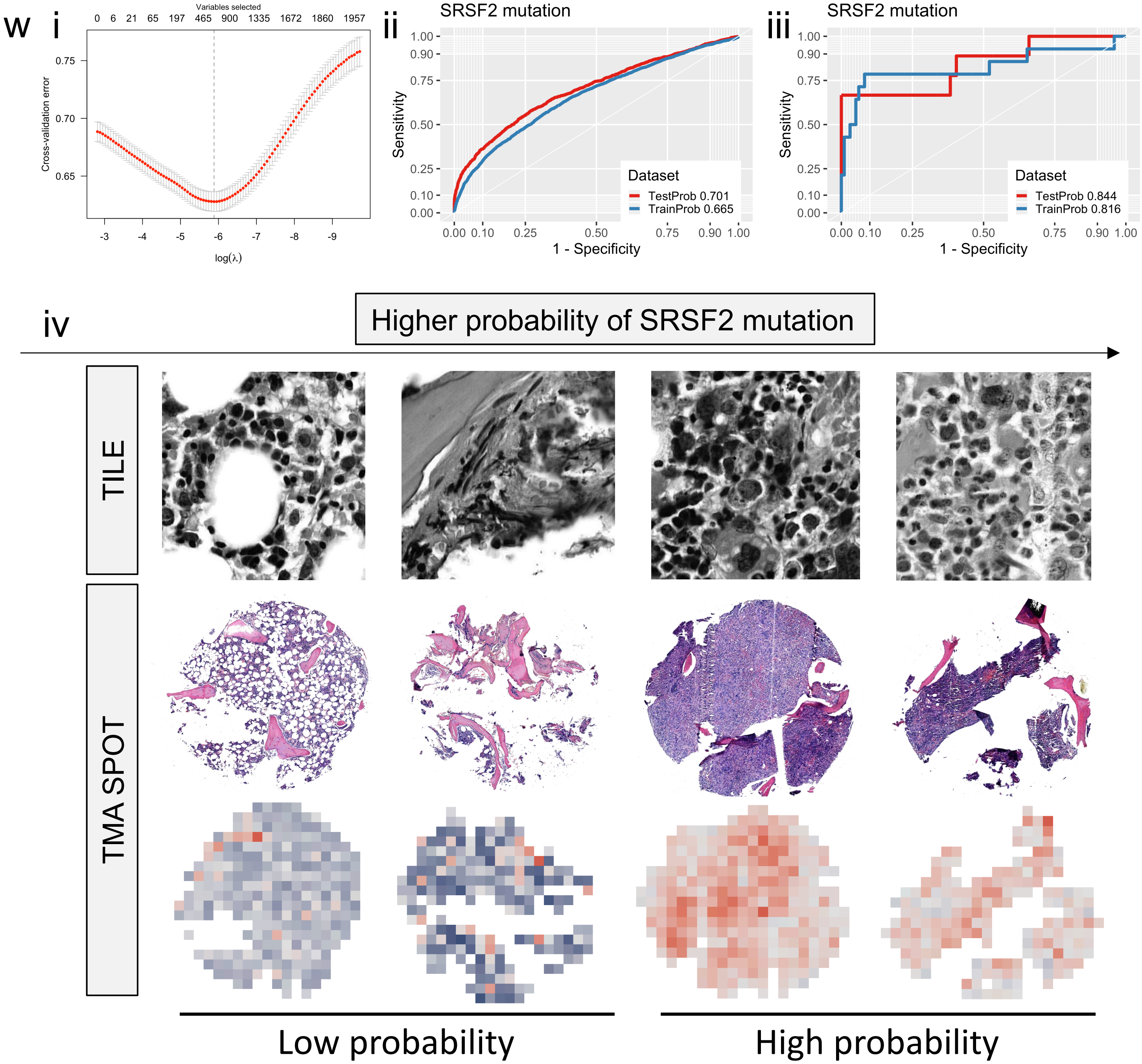


Extended Data Figure 6w.

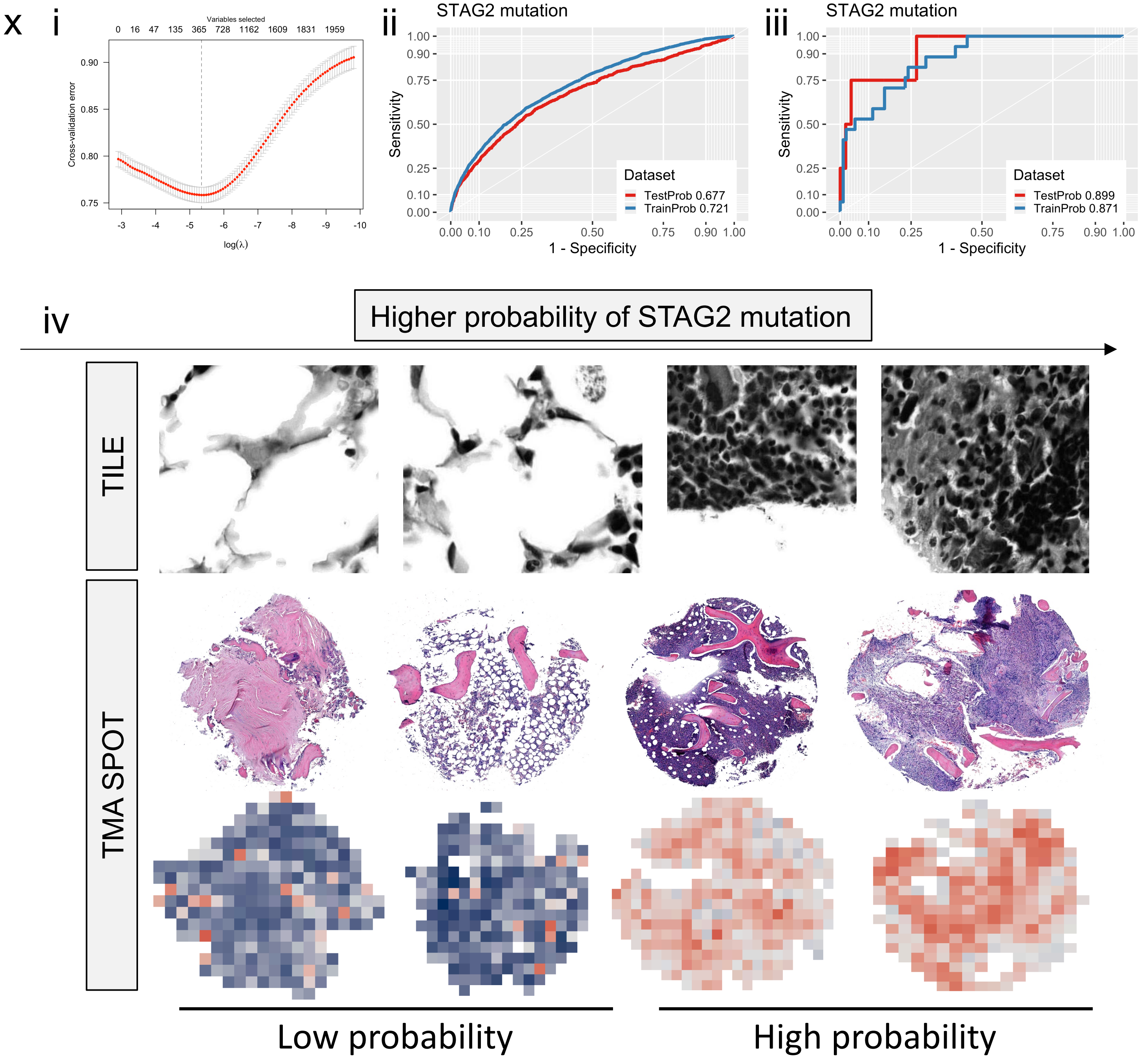


Extended Data Figure 6x.

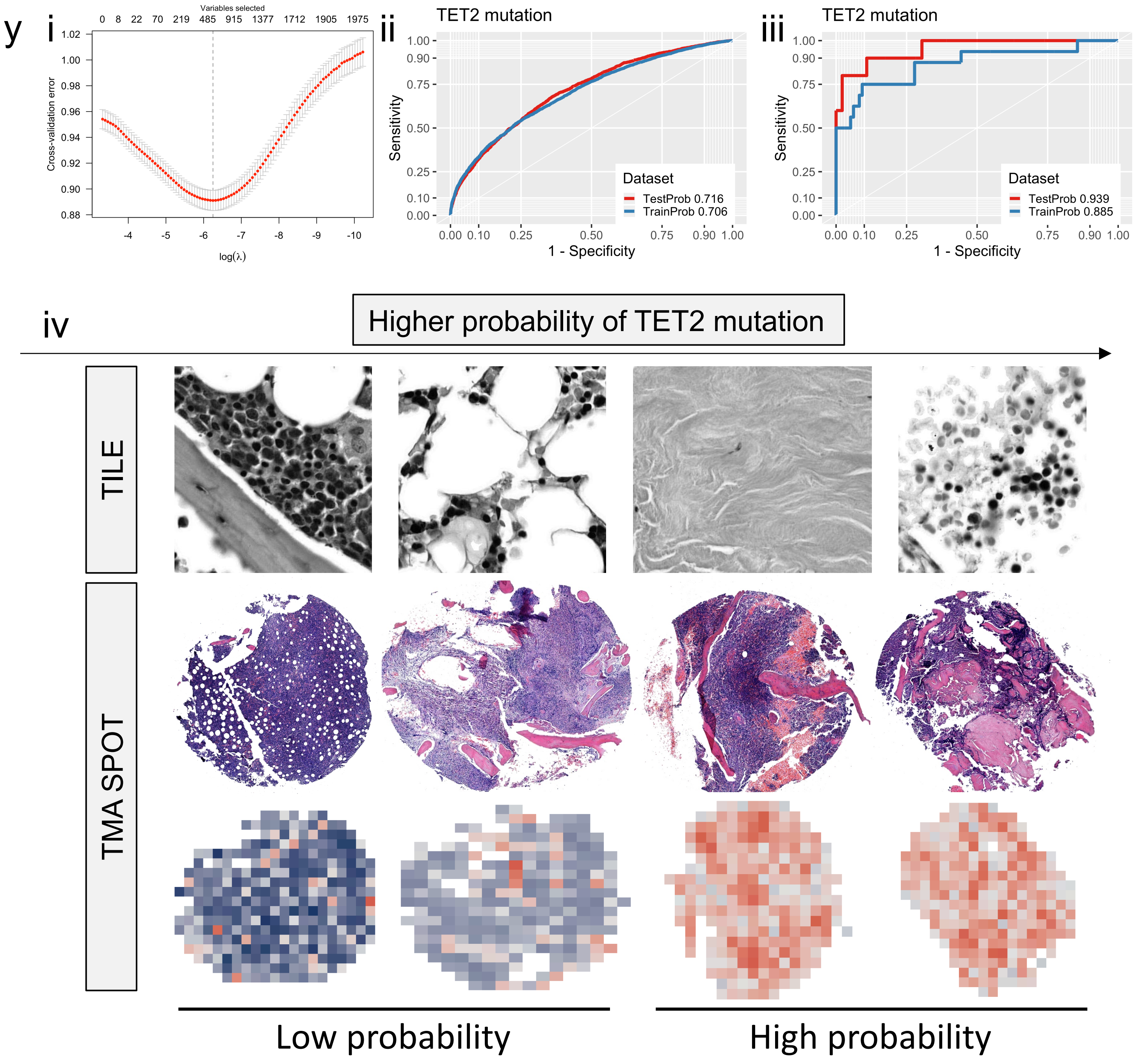


Extended Data Figure 6y.

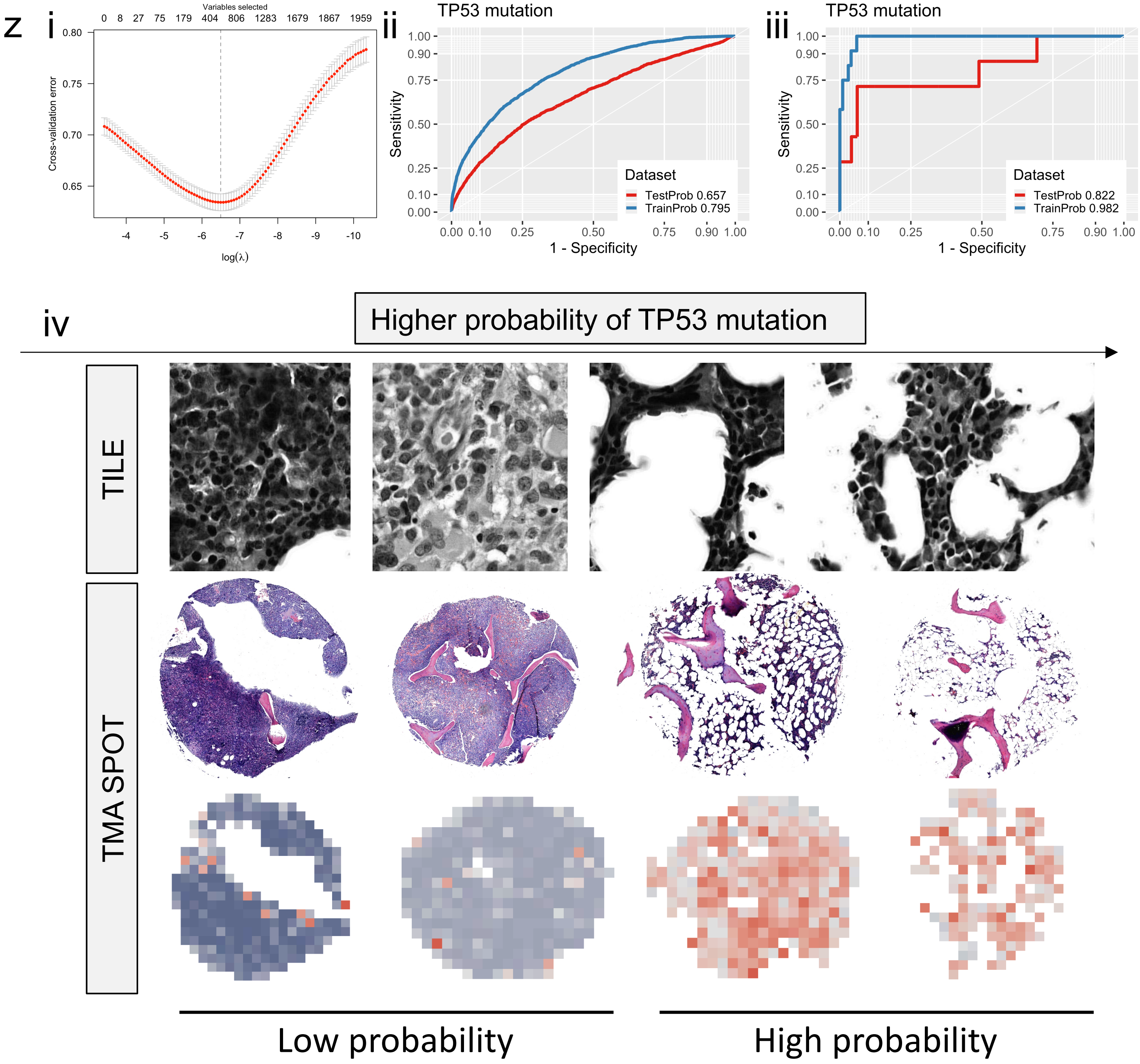


Extended Data Figure 6z.

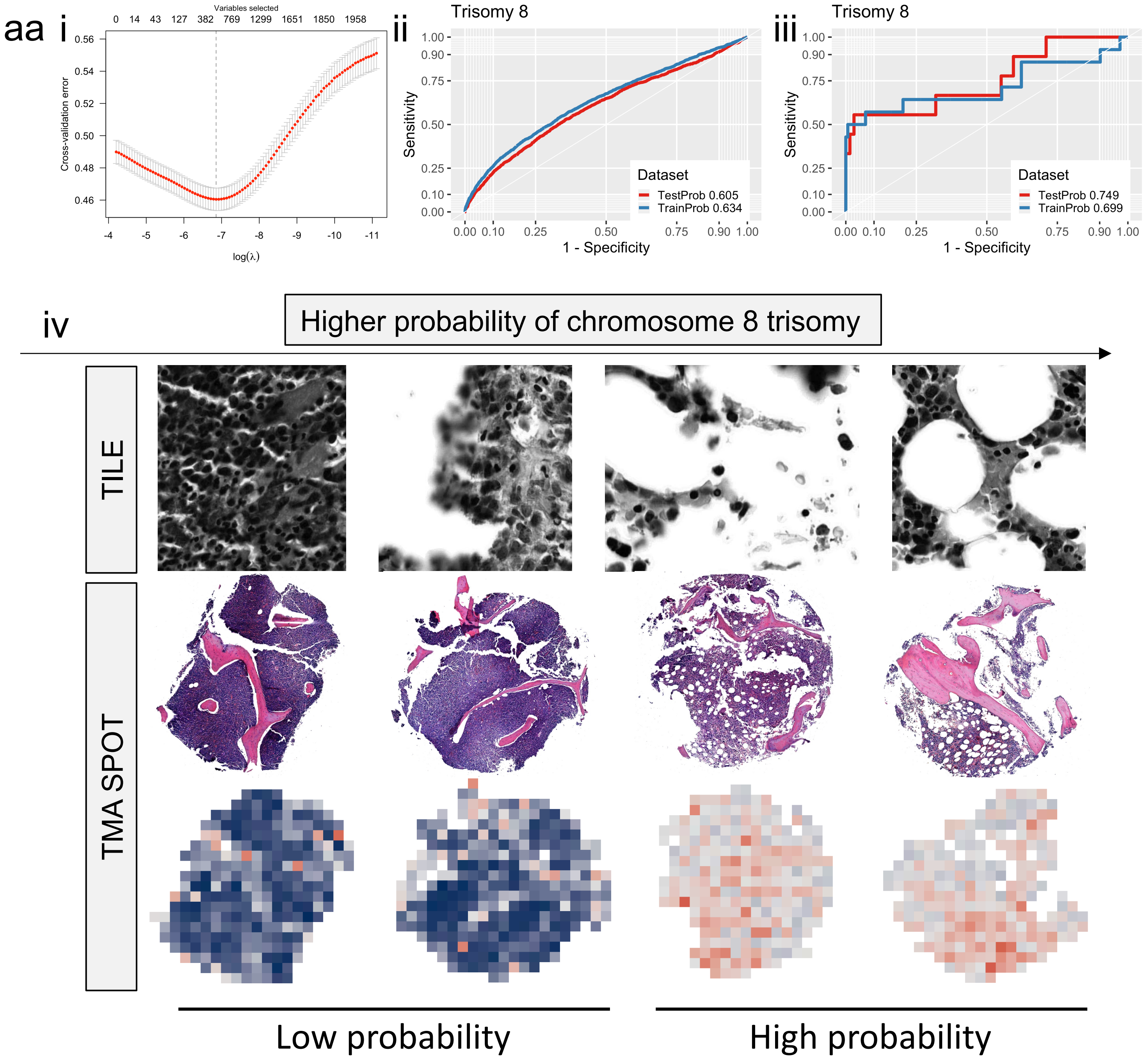


Extended Data Figure 6aa.

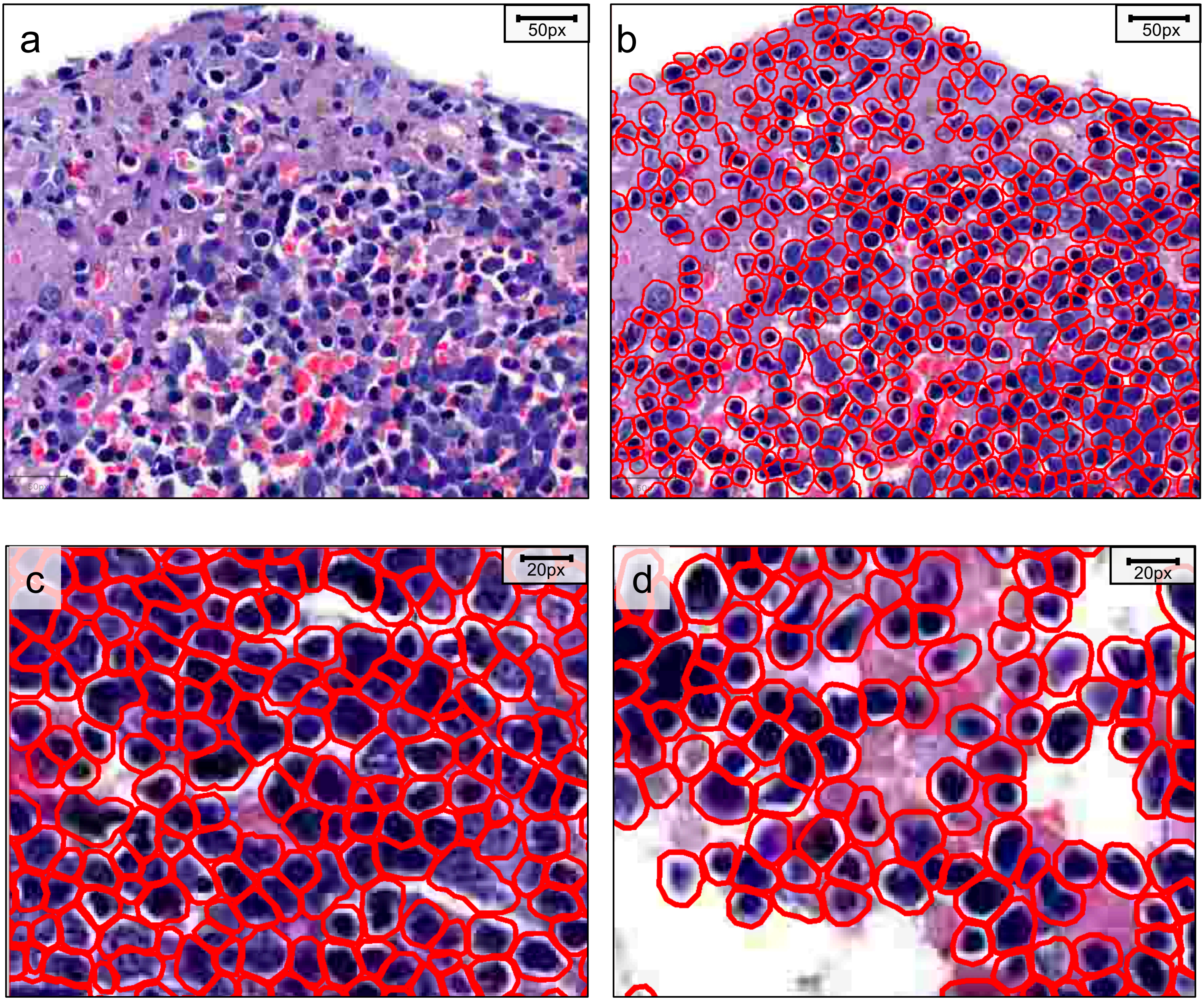


Extended Data Figure 7. (a) Hematoxylin and eosin-stained bone marrow tissue and (b) results from cell segmentation where red-colored boundaries delimit individual cells. Magnified image of cell segmentation in area of (c) high and (d) low cellularity.

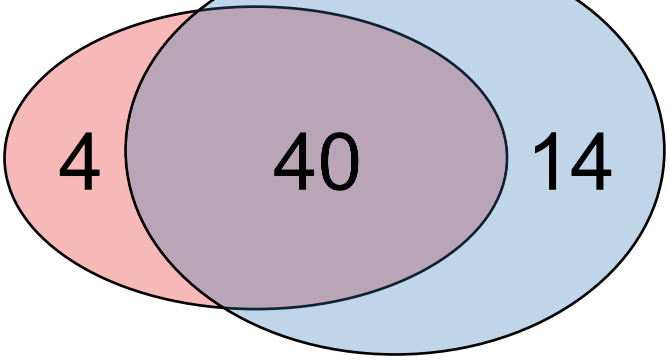


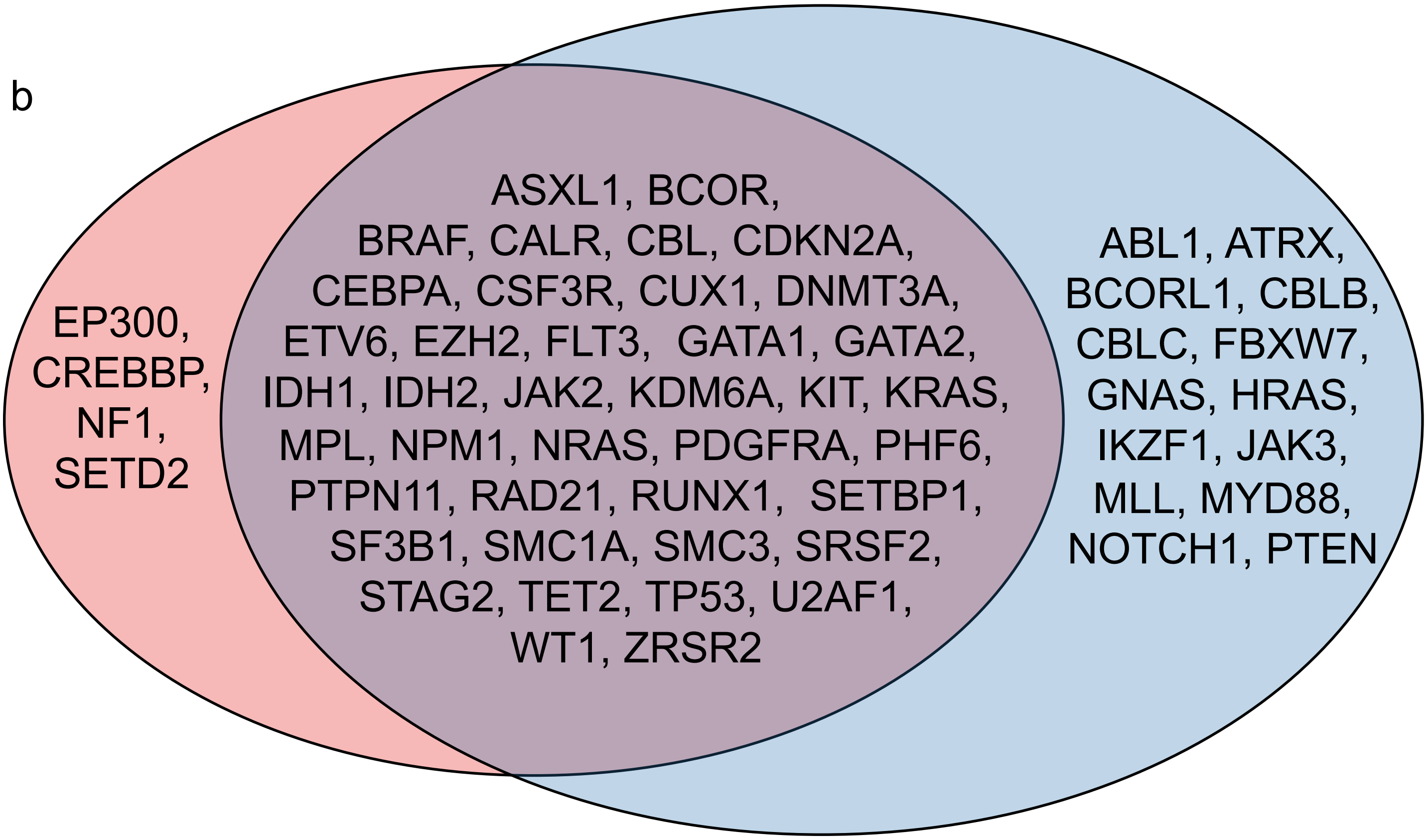


Extended Data Figure 8. Venn diagram visualizing (a) number and (b) gene names included in myeloid amplicon panels used in the study. Gene names on red background are part of the HUSLAB myeloid panel (108 samples) and blue background of the Illumina TruSight panel (30 samples). Genes included in both panels were analyzed in the study.

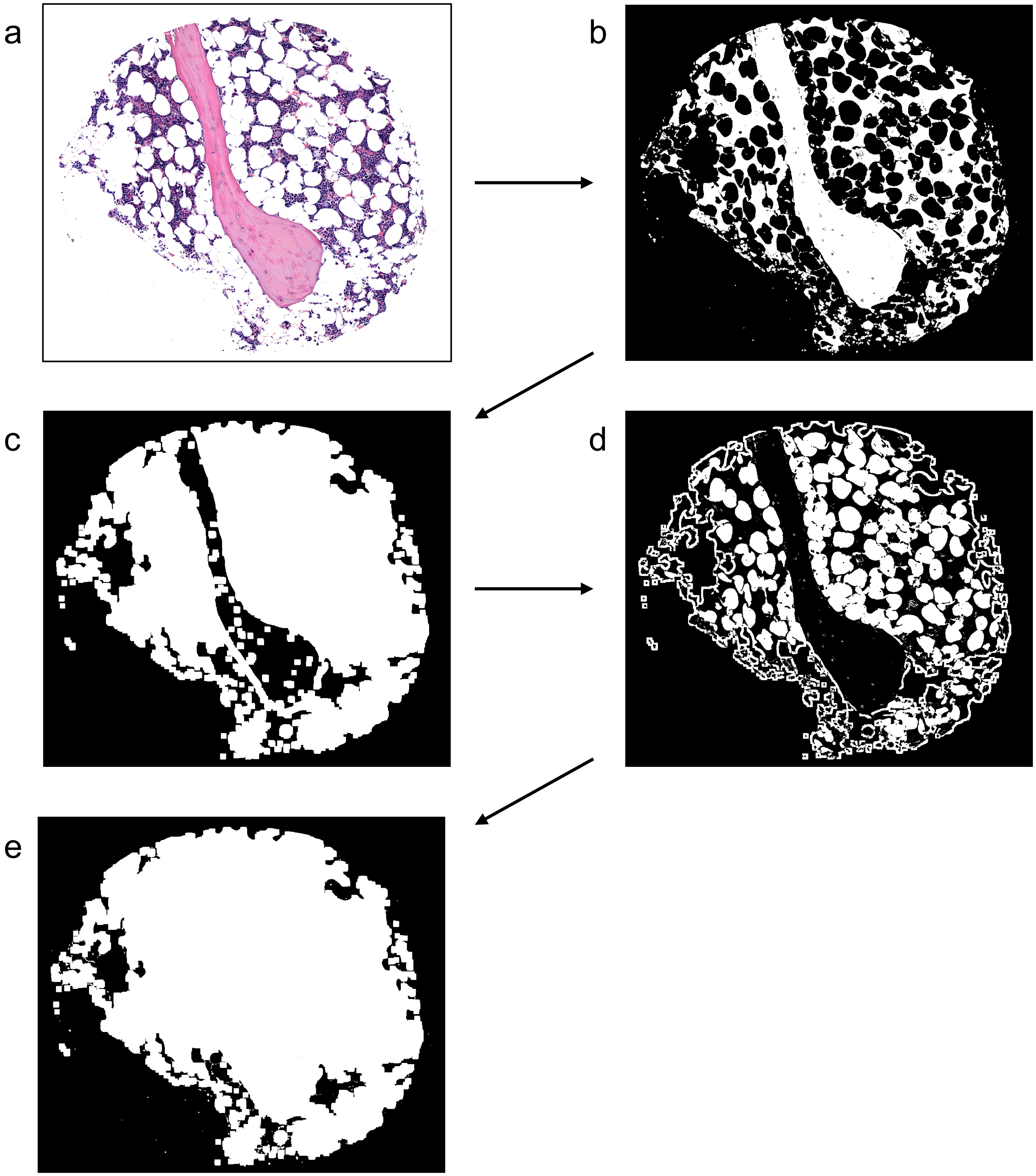


Extended Data Figure 9. Tissue mask creation. (a) H&E-stained tissue microarray core of bone marrow (BM) biopsy. (b) Binary image of figure A. (c) The binary image mask has been dilated and empty holes filled. (d) Filled holes in image C are visualized here with white color. (e) The final image mask consists of combined white areas in images B and D.

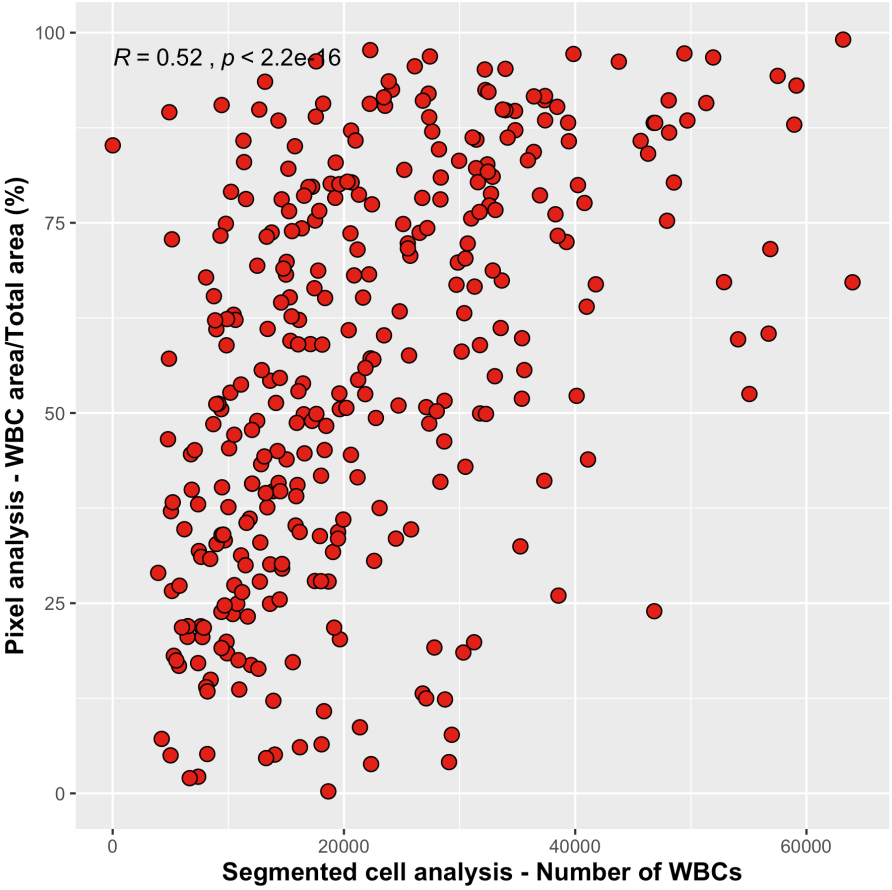


Extended Data Figure 10. Spearman correlation between white blood cell (WBC) area quantitated with pixel classification (y-axis) and WBC cell count defined by cell segmentation (x-axis).
